# Why don’t transcription factors exploit inter-nucleotide correlations in binding motifs? Evolution of information encoding strategies on DNA

**DOI:** 10.64898/2026.11.09.765386

**Authors:** Emmanuel Mekasha, Elia Mascolo, Ivan Erill

**Author notes:** These authors have contributed equally to this work and share first authorship (Emmanuel Mekasha, Elia Mascolo). Corresponding authors: Elia Mascolo and Ivan Erill.

## Abstract

Transcription factors have been shown to predominantly rely on independent contributions from each position of their target sequence motifs, even though sequence motifs can encode information through correlations among nucleotide positions. As a result, the position-weight matrix, a simple modeling approach that assumes independence among positions, has long since become the dominant model of TF-DNA binding. Why evolution would favor this comparatively simple strategy, known as single locus information (*SI*), over a correlation-based one remains unaddressed. Here, we use analytical derivations, numerical approaches and evolutionary simulations to show that *SI* is intrinsically more robust to mutation than correlated (multi-locus) encodings and is therefore indirectly favored under selection for functional transcriptional regulatory networks. At the same time, we show that *SI* smooths the affinity landscape, ensuring the evolvability of regulatory interactions. We find the same marked bias toward *SI* in the genetic code, suggesting that the preference for *SI* is not specific to transcriptional regulation, but may be a general property of biological codes that must reliably store information on DNA in the face of mutational noise.

## 1. Introduction

The binding of molecules to specific locations within nucleotide sequences underpins many fundamental cellular processes. Among these, transcriptional regulation plays a central role by controlling gene expression in response to endogenous and exogenous signals (Ptashne 2005). Transcription factors (TFs) are specialized proteins that regulate the expression of target genes by binding to DNA within cognate regulatory regions, either facilitating or hindering the formation of an open promoter complex (Orphanides and Reinberg 2002). The set of regulatory interactions between TFs and their target genes is commonly referred to as the transcriptional regulatory network (TRN). Most TFs bind to short (5-30 bp) contiguous sequence patterns, or motifs, which they must recognize within a much larger genome sequence. The modeling of TF-binding motifs generated from the alignment of experimentally validated target sites has been the focus of a substantial body of work in bioinformatics (Osada et al. 2004). Early work on TF-binding motifs pioneered the use of position-specific frequency matrices (PSFM) to model TF-binding preferences and search for motif instances in genomic sequences (Staden 1984). This approach rests on the assumption that each position of the motif contributes to TF-binding affinity independently of the others. Under this assumption, a binding-site score is computed as the sum of individual position contributions derived from the observed base frequencies at each position in the binding sites alignment (**Figure 1a-b** and **Figure 1e-f**).

**Figure 1.**
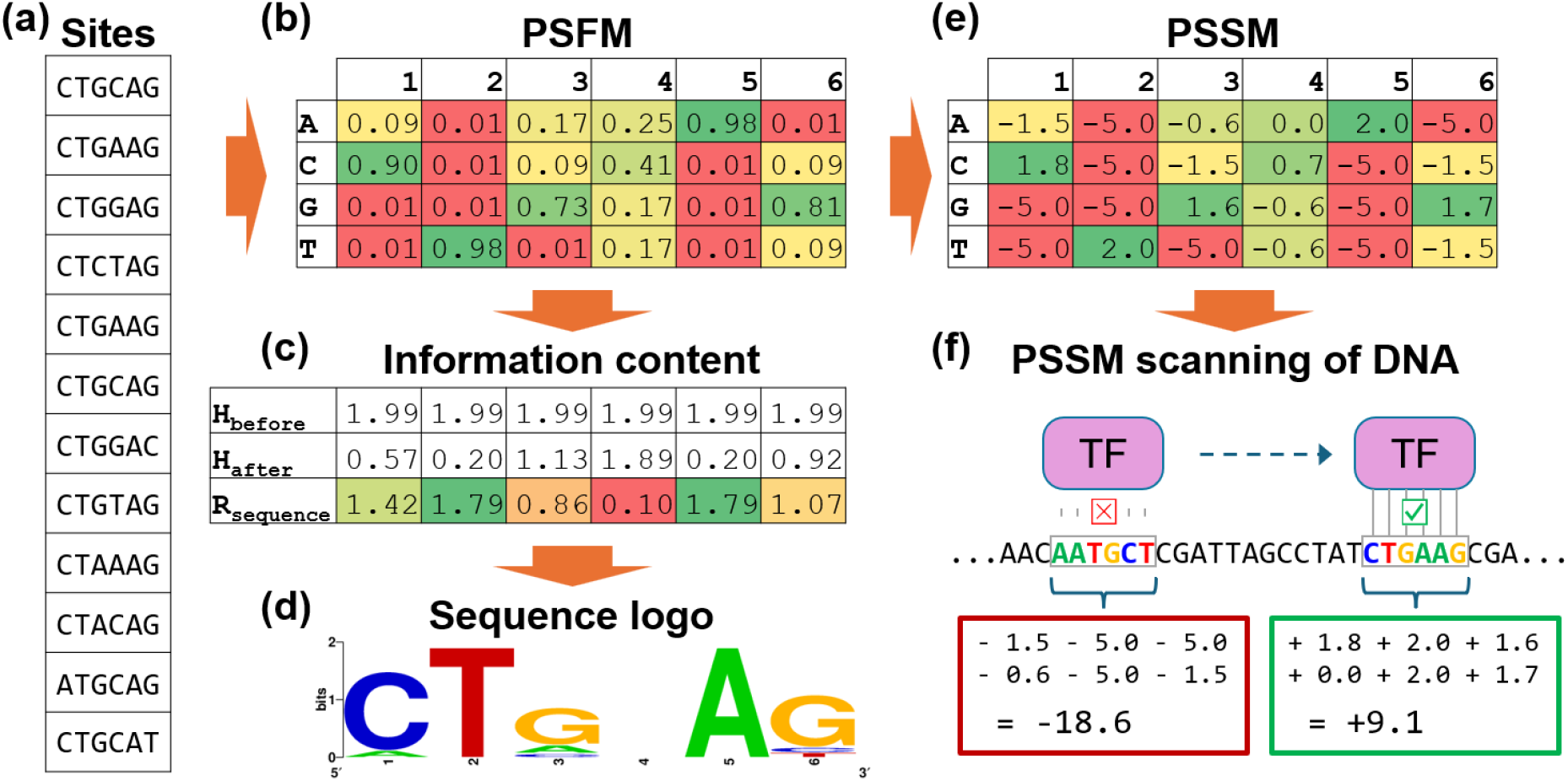
Schematic visualization of TF-binding motif representation. **(a-b)** Illustration of the generation of a position-specific frequency matrix (PSFM), obtained by considering the observed frequency of each nucleotide at each position of aligned TF target sites (experimentally obtained). **(c-d)** The information content can be calculated and displayed through a sequence logo, where the height of each stack of letters at each position represents the total information content at that position (from 0 to a maximum of 2 bits per nucleotide position). **(e-f)** The PSFM can be turned into a position-specific scoring matrix (PSSM), by applying a log-likelihood ratio, that can be used to score each possible *L*-mer (in the illustrated example *L*=6), based on how well it matches the TF binding preferences modeled by the PSSM. A negative score and a positive score are illustrated, representing the prediction of a non-target (−18.6) and a target site (+9.1), respectively.

Despite its simplicity, the PSFM formalism remains the dominant paradigm for modeling TF-DNA interactions. However, the assumption of independence is not mandated by biophysics. In principle, correlations between nucleotide positions can encode additional information about binding affinity, reflecting epistatic interactions among positions due to structural constraints, cooperative contacts, or other context-dependent effects. Motivated by this possibility, numerous studies have proposed motif models that incorporate dependencies between positions, ranging from pairwise to higher-order terms. While such models can capture correlations present in some datasets, their predictive performance has generally improved modestly relative to independence-based models, and in many cases not significantly (Osada et al. 2004; Kulakovskiy et al. 2013; Omidi et al. 2017). Consistent with this, experimental evidence has confirmed that PSFM-based models provide a very good approximation to TF-binding affinity (Benos et al. 2002; Zhao and Stormo 2011; Weirauch et al. 2013; Zuo and Stormo 2014).

These empirical findings raise a fundamental evolutionary question: if correlated encodings are a viable strategy to achieve binding specificity (Le et al. 2018), why do TF-binding motifs predominantly rely on independent positions?

Information theory offers a natural language to formalize this question. The specificity of a TF motif can be viewed as a reduction in sequence uncertainty relative to the genomic background (T. D. Schneider et al. 1986) (**Figure 1**c-d). The *a priori* uncertainty over which base occupies a given position *l* within a genomic fragment of length *L*, is given by Shannon entropy (Shannon 1948):

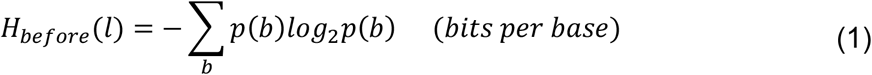

where *p*(*b*) is the genomic frequency of DNA base *b* ∈ {*A, C, G, T*}.

Conditioning on TF binding reduces this uncertainty: the *a posteriori* uncertainty at position *l*, determined by the TF’s base preferences, is

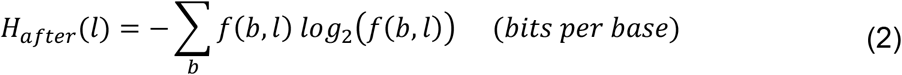

where *f*(*b, l*) is the frequency of base *b* at position *l* of the binding sites.

Following Shannon’s definition of information, we can express the information gained on that position as the decrease in uncertainty:

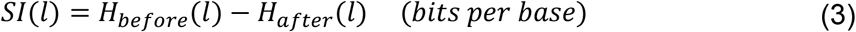

Under the assumption that each position contributes independently to TF-binding, the *single-locus information* (*SI*) for the entire TF-binding motif is simply the sum of the information gained at each position:

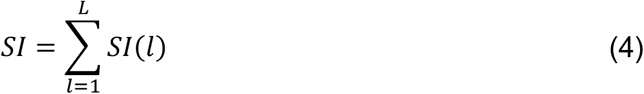

This quantity is also known as *R*_*sequence*_ or *IC* in the literature (T. D. Schneider et al. 1986; Thomas D. Schneider 2010; Stormo 2013) because it is the standard metric for quantifying the information content of a biological sequence, and has been shown to be approximately the amount theoretically required to provide sufficient specificity for molecular recognition of TF target sites (T. D. Schneider et al. 1986; Thomas D. Schneider 2010).

However, single-locus information does not equate to the *total information* of the motif, as sequence patterns can also achieve specificity through correlations among positions. When two positions *X* and *Y* are correlated, the uncertainty at one position can be further reduced based on observing the state of the other. This information gain can be measured (in bits) as *mutual information* (*MI*):

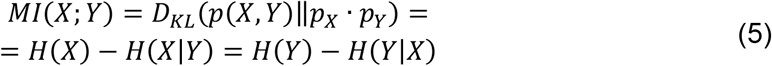

where *D*_*KL*_ is the Kullback-Leibler divergence (Kullback and Leibler 1951), which here measures the deviation (in bits) from statistical independence. *MI* hence measures the information in the alignment that cannot be obtained from the probability distributions at each individual position, but which rather stems from correlations among the positions.

Importantly, *SI* and *MI* represent distinct and complementary sources of information. The total information (*TI*) for a pair of positions can be decomposed exactly into the sum of these two quantities (proof in Appendix A):

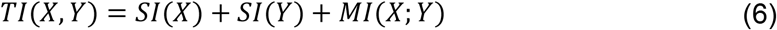

Mutual information can be generalized to capture correlations among an arbitrary number of positions through the quantity known as *total correlation* (Watanabe 1960) (or *multi-information*), which allows us to write the more general decomposition

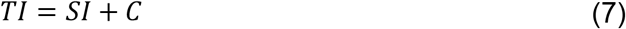

where *TI* is the total information of an *L*-bp motif, *SI* is its overall single-locus information (Eq. 4), and *C* denotes the total correlation (derivation in Appendix A.2).

In practice, however, estimating higher-order correlations requires exponentially large sample sizes. Because TF-binding motifs typically comprise at most a few hundred sites, the analyses of correlation in these systems have been largely restricted to pairwise mutual information. Moreover, whenever present, the correlations contributing to TF-binding affinity are mostly between adjacent positions (Zhao et al. 2009, 2012; Le et al. 2018), and are commonly interpreted as arising from stacking interactions that generate local DNA shape features conducive to TF binding (Zhou et al. 2015). For these reasons, in this work we restrict our informational analysis of TF-binding motifs to pairwise adjacent correlations (see Methods, Section 4.1).

In this work, we investigate whether the structure of the information-encoding strategy itself influences the mutational robustness and evolvability of TF-binding affinity. We compare motifs with identical total information that differ in how that information is encoded: only through independent positional contributions (pure *SI*), only through correlations between positions (pure *MI*) or through mixed strategies (varying percent *SI*). By combining analytical results, numerical approaches, evolutionary simulations and landscape analysis, we ask whether independence could emerge as an evolutionarily favored encoding strategy under mutational noise. Finally, we extend the same informational framework beyond transcriptional regulation by examining the genetic code, asking whether a bias toward *SI* reflects a general principle that influences how biological information is encoded in DNA.

## 2. Results

### 2.1 Asymptotic information loss rates due to mutation

The ability to maintain encoded information in the face of mutation is a fundamental constraint on biological systems (Masel and Trotter 2010; Rozhoňová et al. 2024). A possible explanation for the observed predominance of *SI* over *MI* in TF-binding motifs is therefore an intrinsic advantage at maintaining information when subject to mutations. To examine this possibility, we quantified the intrinsic sensitivity of these two encoding strategies to mutational perturbation. First, we derived analytical expressions for the asymptotic information loss due to random point mutations. We compared *SI* and *MI* motifs under identical initial total information content (*TI*). Specifically, we analyzed motifs of length *L*=2 bp (the minimal system in which *MI* can be defined) and constrained all motifs to have (1) equiprobable bound sequences and (2) an average of 1 bit per position (consistent with the typical information density of TF-binding motifs (Stewart et al. 2012), yielding a total information of 2 bits in each motif). While *MI* is symmetrical over the two positions, *SI* can freely distribute information unequally among them, opening a range of possibilities for the distribution of the 2 bits over two positions. Therefore, we compared three encoding strategies: ***SI* Case A**, where information is distributed in the most unequal way (2 bits at one position and 0 at the other); ***SI* Case B**, where information is distributed as equally as possible (1 bit at each position); and the ***MI*** motif, which encodes the same total information (2 bits) via correlation among the two positions (**Figure 2**).

**Figure 2.**
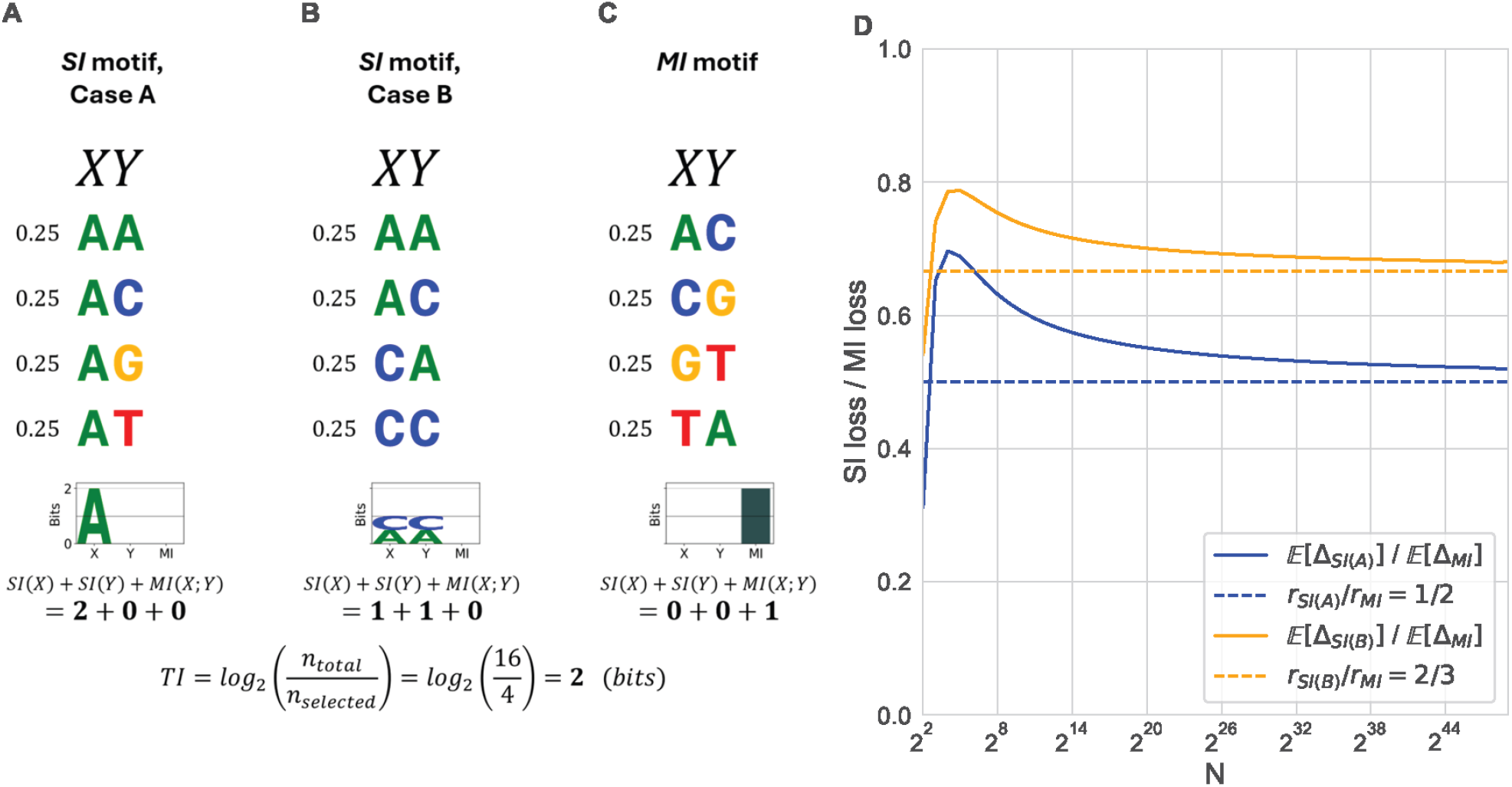
Information encoding in minimal 2-bp motifs and asymptotic information loss. **(A-C)** Examples of 2-bp sequence motifs carrying 2 bits of total information (by selecting 4 out of 16 possible 2-mers; see equation at the bottom), corresponding to an average of 1 bit per position. For each of the three categories, equivalent representatives with the same structure can be obtained by permuting the four letters of the nucleotide alphabet. **(A)** Pure *SI* motif, with maximally unequal distribution of information among positions. **(B)** Pure *SI* motif, with maximally equal distribution of information among positions. **(C)** Pure MI motif, where all information arises from correlations between positions. **(D)** Comparison of the expected information loss after point mutation for finite values of *N*. Due to the equiprobability assumption (assumption (1) in Section 2.1), the frequency of each of the four selected sites of the binding motif is simply *N*/4. As *N* grows, the information loss ratios approach the asymptotic limits of 1/2 and 2/3 for *SI* Case A and B, respectively.

A single point mutation in one of the *N* aligned binding sites that compose the motif will alter frequencies of the underlying distribution *P* by an order *O*(1/*N*). Thus, to obtain the derivative of information with respect to motif perturbation *ε*, namely the initial asymptotic information loss rate

*r*_*I*_, we calculated the limit

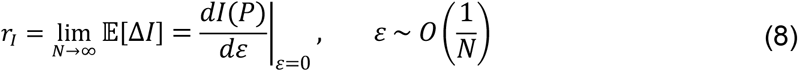

and found the following loss rates (in bits per mutation):

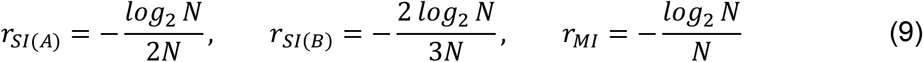

Therefore, *SI* Case A loses information at half the asymptotic rate of the *MI* encoding (1:2 ratio), whereas *SI* Case B shows an intermediate loss rate corresponding to a 2:3 ratio relative to *MI*:

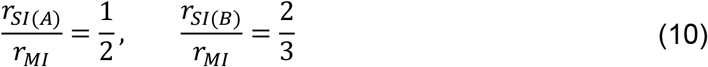

These results show that, under minimal assumptions, *MI* is the most fragile encoding strategy, whereas the mutational robustness of *SI* depends on the specific encoding used (see Appendix B for full derivations).

### 2.2 Lower information loss leads to higher mutational robustness in *SI*-encoded TRNs

The analytical results presented above rely on assumptions introduced for mathematical tractability and define the information loss rate due to infinitesimal changes in an idealized motif. To test whether the predicted differences between *SI* and *MI* extend beyond these idealized cases, we next adopted a numerical approach based on randomly sampled motifs with a given total information content. We applied random mutations to samples of 100 2-bp motifs encoding 1 bit of information either via *SI* or via *MI*, and monitored information loss. We found that *MI*-based motifs lose information substantially faster than *SI*-based ones (**Figure 3**A). The observed information loss rates after the first mutation event approximate a 2:3 ratio (**Figure 3**A - inset), in agreement with the analytical results for *SI* Case B. This alignment of experimental results with *SI* Case B reflects the fact that motifs with a roughly equal distribution of information across positions, as in *SI* Case B, vastly outnumber those with information concentrated in a single position (**Figure S1**).

**Figure 3.**
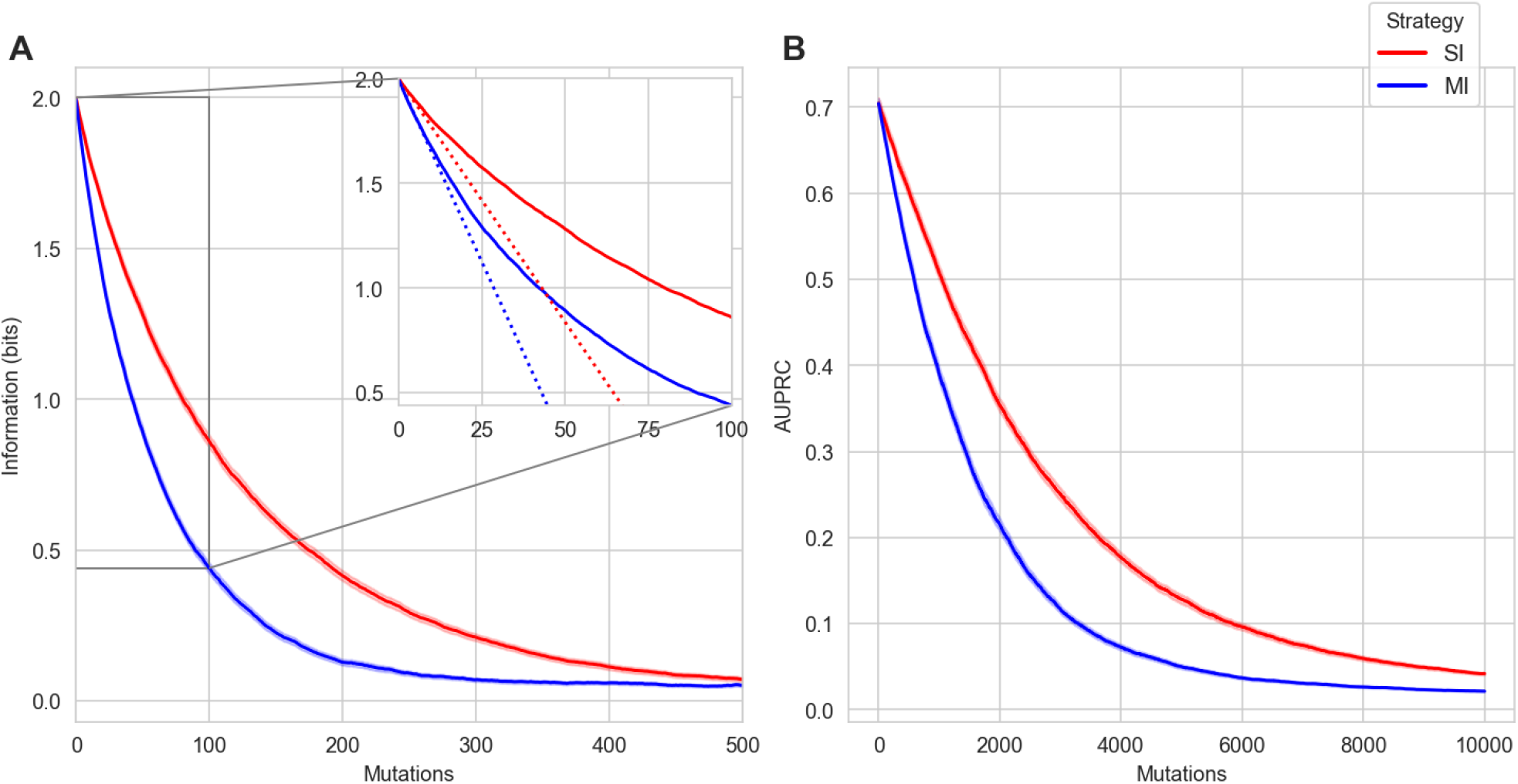
Information loss and AUPRC decay under mutation for MI and SI-encoded TRNs. (**A**) Total information content of *SI*-based and *MI*-based motifs as random mutations on the binding sites accumulate in 2 bp-long sequence motifs. The average (solid line) and the standard error of the mean (shade) are plotted for 100 randomly generated motifs for each population. The empirical slopes for *SI* and *MI* information loss after one mutation (inset: red and blue dotted lines) are, respectively, 0.0233 bits/mutation and 0.0348 bits/mutation. (**B**) AUPRC for genome scans of *SI*- and *MI*-based TRNs as random mutations accumulate in 8192 bp-long genomes. TRNs are composed of 128 target sites, which are 12 bp-long, and encode 6 bits of information. The average (solid line) and the standard error of the mean (shade) are plotted for 100 randomly generated *SI*-based TRNs (red) and 100 randomly generated *MI*-based TRNs (blue).

Extending this analysis to motifs encoding information through mixed contributions of *SI* and *MI* revealed that mutational robustness scales with the fraction of information encoded via *SI*, with motifs relying exclusively on *SI* showing the slowest information decay (**Figure S2**). This pattern persists across different information densities (**Figure S3**), and when generalizing from motifs of length two to motifs of arbitrary length (**Figure S4**).

The loss of the information encoded in TF-binding sites is a challenge for transcriptional regulatory systems because it decreases the ability of a TF to discriminate between its target binding sites and all other genomic locations and, hence, maintain the integrity of the TRN. Treating the TF as a classifier that must distinguish its target sites from the other genomic locations based on binding affinity, we can assess TRN integrity using the area under the precision-recall curve (AUPRC) following a genome scan for TF-binding sites (Kılıç and Erill 2016; Zhang et al. 2024). Therefore, we sampled 12-bp long motifs encoding information preferentially through either *SI* or *MI*, and we embedded each motif’s target binding sites in a random genome sequence. As we introduced random mutations, we monitored the decrease in AUPRC resulting from scanning the genome with the TF model defined by the original set of TF-binding sites. As expected, *SI*-based TRNs showed higher robustness to mutations, measured as a slower loss in classification performance as mutations accumulate (**Figure 3**B), regardless of the length or the information density of the motif (**Figure S5**; **Figure S6**).

### 2.3 Mutational robustness promotes the evolution of *SI*-based TRNs

The higher mutational robustness of *SI*-encoded networks suggests that they may have an evolutionary advantage under stabilizing selection, since the expected fitness of their offspring will be higher. To test this possibility, we devised evolutionary simulations for organisms encoding TF-binding models and their target sites on a randomly initialized genome background, using the TRN integrity (AUPRC) as a proxy for fitness. We applied this framework to conduct *in silico* competition experiments between evolved organisms that encode their TRN either preferentially through *SI* or preferentially through *MI*. Both sets of organisms were equally accurate at the beginning of the simulation (same fitness), but *SI* strategists systematically outcompeted *MI* strategists and reached fixation in the vast majority of the simulations (**Figure 4**A), despite the small population size.

**Figure 4.**
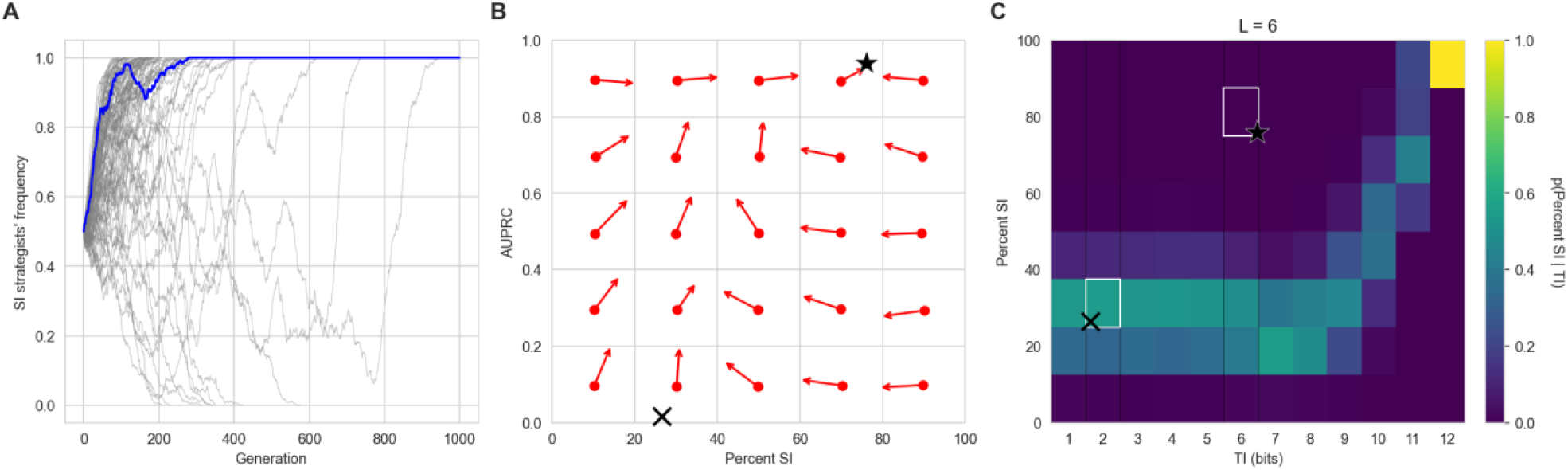
**(A)** Competition experiment involving two TRN strategies: (1) *SI*-strategists encoding 70% of the motif information as *SI* and 30% as *MI*; (2) *MI*-strategists encoding 70% of the motif information as *MI* and 30% as *SI*. At generation 0, both strategies are at equal frequency (0.5) in the initial population, containing 1000 clonal networks per strategy. In all 100 independent simulations, one of the two strategies reaches fixation within 1000 generations. The vertical axis reports the frequency of *SI*-strategists in the population. **(B)** 2D phase space for the evolution of TRNs, parametrized with percent *SI* (horizontal axis) and TRN AUPRC (vertical axis). The phase space is discretized into 25 cells (5 × 5 grid). Clonal populations of organisms centered on each cell evolve until the population’s centroid reaches the cell boundary. All the organisms are simulated using the default parameters (*l*_*genome*_ = 2048 *bp, l*_*bs*_ = 6 *bp, n*_*bs*_ = 32). The average displacement vector for 10 independent experiments is shown as a red arrow in each cell. The steady-state attractor at mutation-selection balance is represented by a star. The mutational steady state is represented by a cross. **(C)** Probability distribution of percent *SI* (vertical axis) conditioned on total information (*TI*) in 6 bp-long motifs. Attractors from panel (B) are mapped onto the probability space and represented with the same symbols.

**Figure 5.**
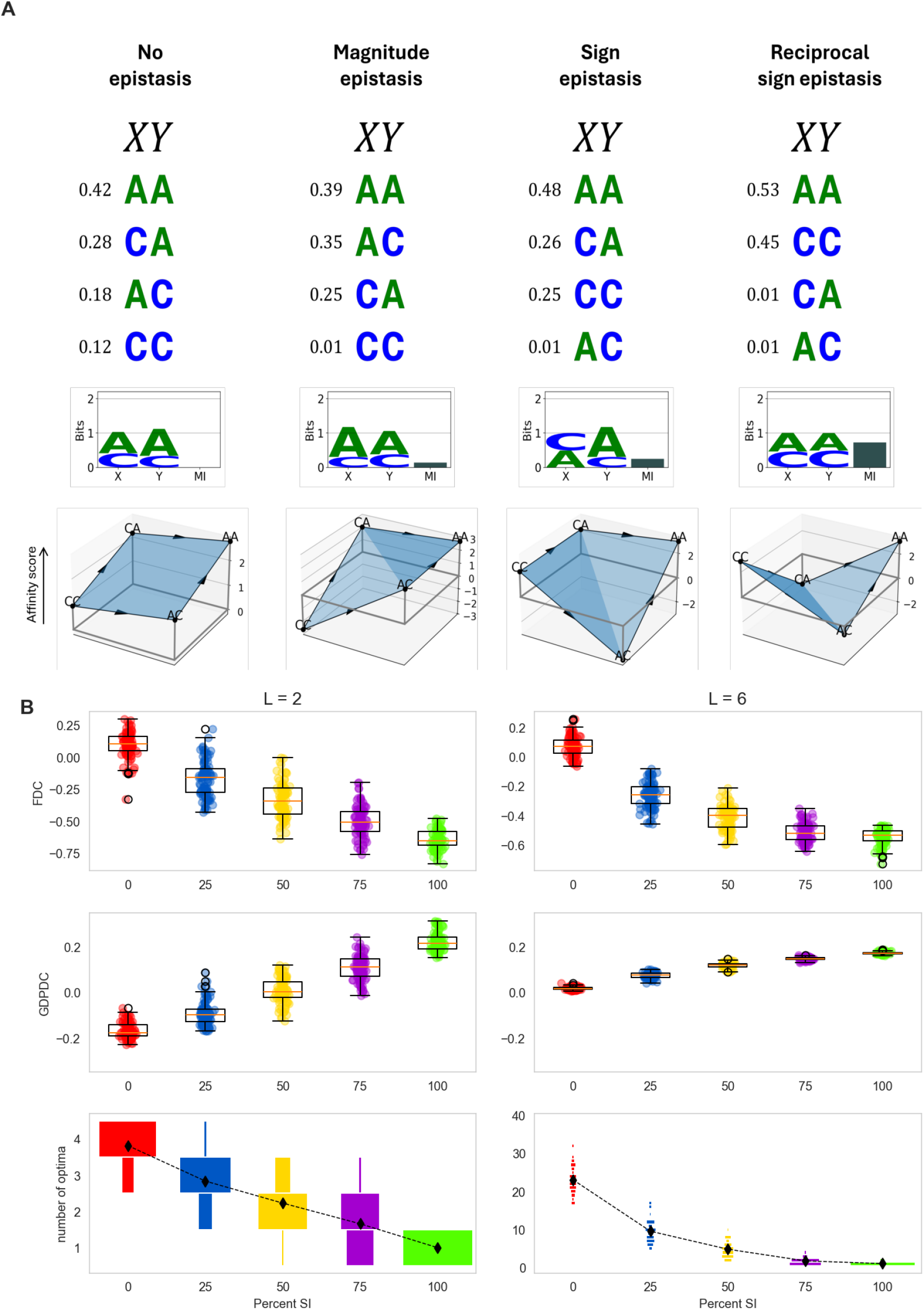
**(A)** Minimal example showing simple (*L*=2) motifs where the same four 2-mers (out of 16) are selected with different probabilities. The corresponding phenotype landscapes are shown, where the reported phenotype is the TF binding affinity score for each sequence, calculated with the standard log-likelihood ratio (Hertz and Stormo 1999). The log-likelihood ratio is calculated using the true joint probability distribution relative to the background, which reduces to the canonical PSSM score only for pure *SI* motifs (Methods, Section 4.4). In the motif representation, the *L*-mers are ranked by probability (top to bottom), and the non-represented *L*-mers are assumed to not be recognized (probability≈0). To keep the landscape representation low-dimensional, recognized sequences were chosen so that they involve only two letters (A and C). In all four cases, we consider the effect of a C→A mutation at each position (*X* and *Y*). **(I) No epistasis**. Purely *SI*-based motifs entail a flat (linear), non-epistatic phenotype landscape (see also Appendix D), where the contributions to the phenotype from each position are additive. **(II) Magnitude epistasis**. The phenotype landscape departs from a linear (additive) one, because the effect of C→A mutations at both positions of the initial CC site is less than the sum of the effects of the two individual C→A mutations. **(III) Sign epistasis**. The effect on binding affinity of a C→A mutation at position *X* is negative when *Y*=C, but positive when *Y*=A. Therefore, *Y* has a sign-epistatic effect on *X*. **(IV) Reciprocal sign epistasis**. Each position has a sign-epistatic effect on the other. **(B)** Global characterization of the TF-DNA binding affinity landscape for 2-bp and 6-bp TF-binding motifs that encode information through different strategies, ranging from 0% *SI* (100% *MI*) up to 100% *SI* (0% *MI*). For each strategy, 100 random motifs are sampled (see Methods, Section 4.2), and the resulting affinity landscape is analyzed by calculating the fitness-distance correlation (FDC) metric, where phenotype (affinity) replaces fitness, the genotype distance-phenotype distance correlation (GDPDC), and the number of optima (peaks) in the landscape.

Competition experiments indicate that the impact of mutations accumulating over generations is greater in *MI*-based TRNs, leading to indirect selection for *SI*-based regulatory codes. Therefore, we hypothesized that transcriptional regulatory systems could evolve *SI*-based TF-binding affinity codes due to indirect selection for robustness, even in the absence of any direct fitness advantage, mutation biases or constraints favoring *SI* over *MI*. To test this hypothesis, we characterized the global evolutionary trajectories of freely evolving regulatory systems. We considered a discretized 2D phase space for TRNs, with low-to-high percent *SI* (see Methods 4.1) on one axis and low-to-high TRN functionality, quantified by the AUPRC, on the other. For each represented combination of percent *SI* and TRN functionality, we estimated the expected evolutionary change under selection for AUPRC, and represented it as a displacement vector. The resulting vector field demonstrates that, as DNA accumulates information and the TRN progresses toward functionality (greater AUPRC), the regulatory system experiences indirect selection toward higher *SI*, due to the ability of *SI* to preserve the already evolved regulatory interactions in the face of mutations (**Figure 4**B).

The phase diagram shows a steady-state attractor at ~75% *SI* (marked with a star), with TRNs operating near perfect classification (AUPRC≈0.95). This attractor lies far from the mutational steady state in the absence of selection at ~25% *SI* (marked with an X), which instead favors *MI* simply because the space of motifs that encode information predominantly through *MI* is vastly larger than that of motifs relying primarily on *SI*. To quantify this representational bias toward *MI*, we analyzed the distribution of percent *SI* as a function of the total information (*TI*) in TF-binding motifs by uniformly sampling motifs encoding different amounts of *TI* and computing their percent *SI*. As shown in **Figure 4**C and **Figure S7**, *SI*-encoded motifs are extremely unlikely at low and intermediate information densities, and only become dominant at information densities that strictly require *SI* (i.e., which cannot be reached through *MI* alone), near the maximum density of 2 bits per position. As expected, the steady-state in the absence of selection (*X*) identified in **Figure 4**B maps to the highest probability density region in **Figure 4**C, whereas the steady-state attractor at selection-mutation balance (star) lies in a low probability density region, indicating that it can arise only through selection. Overall, these results demonstrate that, at the information densities typical of TF-binding motifs (~1 bit per position (Stewart et al. 2012)), predominantly *SI*-encoded sequence motifs are unlikely to arise by chance, and suggest that the evolution of *SI*-encoded regulatory networks is driven by indirect selection for mutational robustness, in the face of a representational bias that rather favors *MI*-rich motifs.

The specific value of the mutation-selection steady state reported in Figure 4B (~75% *SI*) is the result of the simulation parameters used in this study, including a mutation rate set well above biologically realistic values for computational tractability, rather than a quantitative prediction of the equilibrium percent *SI* expected in any specific biological system. Different parameter choices will shift this attractor location without changing the qualitative direction of the effect. For example, for prokaryotic TF-binding motifs the relevant parameters are substantially different, including a much bigger population size (which increases selection efficacy) and a much lower mutation rate (which reduces the pull of mutational/representational biases). Consistent with this qualitative prediction, empirically measured percent *SI* values for experimentally-validated prokaryotic TF-binding motifs are substantially higher (~96%; **Figure S8**) than those observed in our simulations.

### 2.4 SI enhances the navigability of the affinity landscape

We provided evidence that *SI*-based TRNs can result from indirect selection for mutational robustness, a mechanism that is relevant when preserving already evolved regulatory interactions is advantageous, that is, under stabilizing selection. TRNs, however, can also be intermittently subject to directional selection in response to environmental changes, favoring small adjustments or even a complete network rewiring (Galardini et al. 2015; Johnson 2017; Adhikari et al. 2021). Such processes may involve the loss, acquisition or fine-tuning of regulatory interactions and, therefore, require that the TF-binding affinity of certain sites be modulated, and not just preserved.

The evolvability of a TRN is hence predicated on how readily different affinity values can be reached through changes in the DNA sequences, with the possible mutational changes and their phenotypic effects defining a binding affinity landscape (Aguilar-Rodríguez et al. 2017; Fragata et al. 2019). The predominance of *SI* in naturally occurring TF-binding motifs therefore raises the question of whether *SI*-based encoding, while conferring mutational robustness under stabilizing selection, is also compatible with efficient navigation of the affinity landscape under directional selection. To address this, we compared the navigability of TF-binding affinity landscapes associated with *SI*- and *MI*-encoded motifs using three complementary metrics. The *fitness-distance correlation* (FDC) measures the correlation between affinity and the sequence with maximum affinity. The *genotype distance-phenotype distance correlation* (GDPDC) measures the correlation between genotype and phenotype distances, across all possible pairs of genotypes. Lastly, the *number of local optima* is used as a proxy for the ruggedness of an adaptive landscape, with fewer local optima denoting a more navigable landscape.

Our results for 2-bp and 6-bp motifs show conclusively that *SI*-encoded motifs generate more navigable landscapes when compared to *MI*-encoded motifs (**Figure *5***). FDC coefficients shift from positive to markedly negative as we increase the percentage of information encoded through *SI* (**Figure *5***B), indicating that TF-binding affinity decreases with increasing genetic distance from the best binding sequence in *SI*-based motifs, but not in *MI*-based ones. Consistent with this finding, GDPDC coefficients increase with percent *SI* (**Figure *5***B), indicating the prevalence of available monotonic paths in the affinity landscapes of *SI*-based motifs, but not in *MI*-based ones. Moreover, the number of optima decreases as the percentage of *SI* increases (**Figure *5***B), indicating that *SI*-encoded motifs generate smoother affinity landscapes. Together, these results suggest that encoding information using *SI* not only increases the mutational robustness of embedded TRNs, but is also compatible with efficient tuning and navigation of TF-binding affinities, consistent with a high potential for TRN evolvability.

### 2.5 Mutational robustness correlates with percent *SI* in the genetic code

Our results indicate that encoding genomic information preferentially through *SI* increases mutational robustness while remaining compatible with, and potentially even facilitating, the tunability and evolvability of regulatory interactions. This advantage is due to intrinsic properties of *SI* and *MI*, rather than the molecular details of TF-DNA recognition. Therefore, we expect it to hold true for other biological codes outside the context of transcriptional regulatory systems. The genetic code is the most extensively characterized code used by organisms to encode information in DNA sequences. From the perspective of information theory, it can be formally conceptualized as a set of 21 sequence motifs encoding 21 messages: the 20 canonical amino acids and the stop signal (**Figure 6**A). To explore the generality of *SI*-preference in the encoding of biological information, we analyzed the standard genetic code under the framework of percent *SI*. We quantified the percentage of *SI* used to encode each message, as well as the overall percent *SI* of the code (see Methods Section 4.1). To quantify mutational robustness, we calculate the probability of incorporating the correct amino acid after a codon undergoes a random point mutation (µ-robustness in (Maeshiro and Kimura 1998)). We compared the standard genetic code to 1,000 randomized codes that preserve its degeneracy structure, that is, the number of synonymous codons assigned to each message, while randomly permuting codon-message assignments. We found that the standard genetic code stands out not only in terms of mutational robustness, which is one of its well-known features (Freeland and Hurst 1998; Maeshiro and Kimura 1998; Goodarzi et al. 2004), but also in terms of percent *SI* (97.39%). Unlike the standard genetic code, random codes display substantial levels of *MI* in most sequence motifs (**Figure 6**B-C). The value of percent *SI* for the standard genetic code is consistent with the one estimated for experimentally-validated prokaryotic TF-binding motifs (96.37%, SEM=0.60 percentage points; Figure S8). Importantly, we also found that, as hypothesized, percent *SI* and code robustness are strongly correlated in randomized genetic codes (**Figure 6**B).

**Figure 6.**
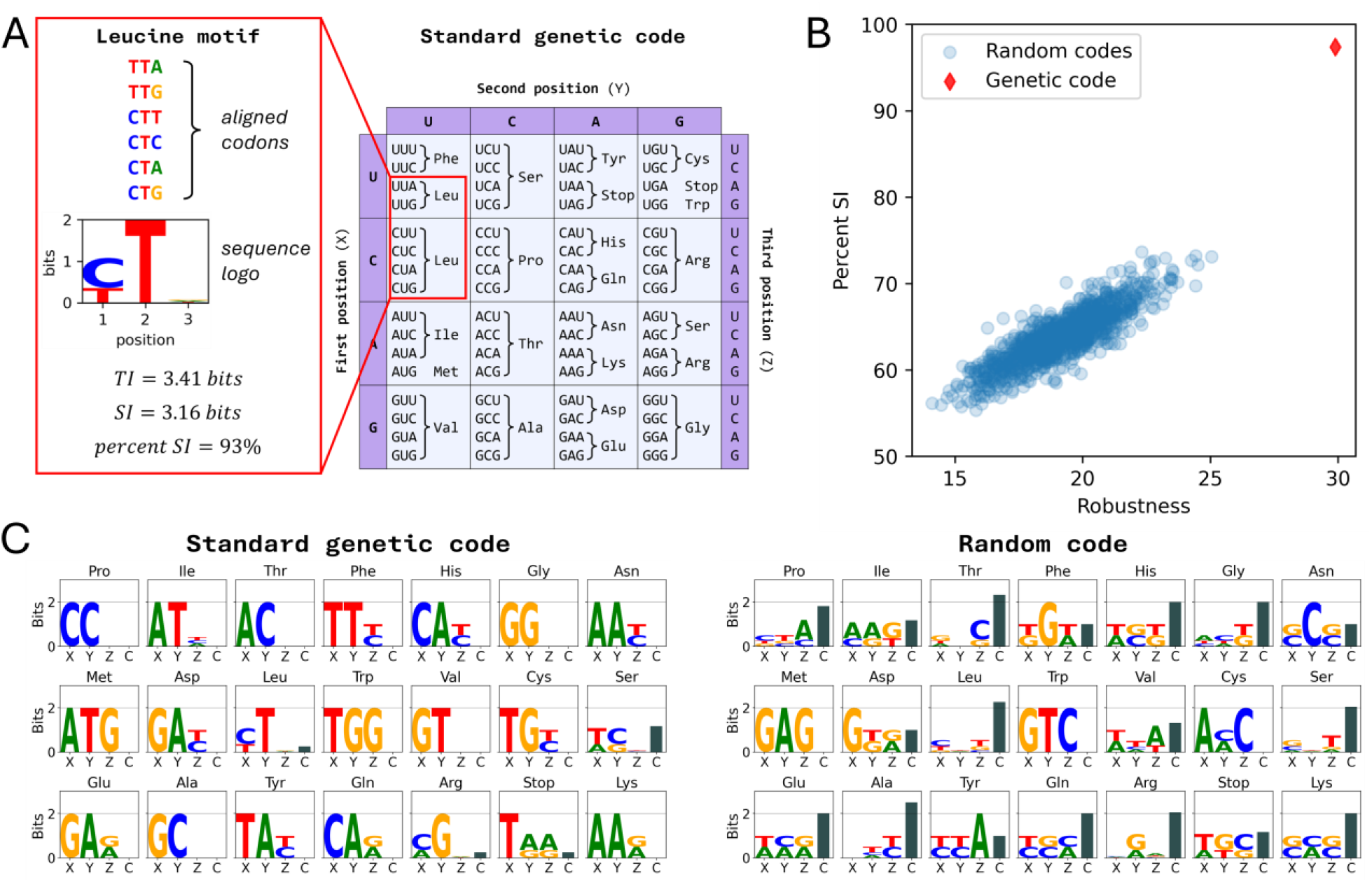
**(A)** The synonymous codons that code for Leucine are shown aligned, and a sequence logo is obtained, showing the *SI* of each of the three motif positions as the height of each stack of letters *(T. D. Schneider and Stephens 1990)*. The total information (*TI*), the overall single-locus information (*SI*) and the percent *SI* for the Leucine-encoding motif are reported. **(B)** The percent *SI* over all the 21 motifs (20 encoding for an amino acid and one encoding for the stop signal) is reported for both the genetic code and for 1000 random codes (random codon-signal assignments that respect the number of codons per signal found in the genetic code). For each code, mutational robustness is estimated as the percentage of single-nucleotide substitutions that result in a mistranslation. **(C)** The sequence logos for the 21 messages as encoded in the standard genetic code are shown on the left. Each sequence logo includes an extra bar quantifying (in bits) the information encoded via correlations among positions (the total correlation, *C*). On the right, the same representation is applied to a randomly generated “random code”. Amino acid motifs in the standard genetic code display a high proportion of *SI*, which is extremely unlikely based on random sampling (**Figure S9**, Methods Section 4.2).

## 3. Discussion

### 3.1 Robustness as a property of the coding structure

TF-binding motifs are known to encode most of the information through independent contributions from each nucleotide, with limited dependencies between them. This observation motivated us to investigate whether this encoding strategy could be an evolved feature that grants mutational robustness. We quantified robustness based on the expected loss of information due to mutation and found that *SI* yields lower information loss rates than *MI*, both in analytically tractable idealized cases and under general conditions evaluated numerically. This formulation of robustness differs conceptually from the classical definition based on neutral network theory, which quantifies the fraction of single-mutant neighbors of a given genotype that preserve its phenotype (thus estimating a phenotype loss probability). Robustness in the classical framework is hence a local property of a specific genotype-phenotype pair, and the phenotype is treated as a discrete trait that is either lost or retained (van Nimwegen et al. 1999; Wagner 2005). In contrast, by measuring expected information loss (in bits per mutation), robustness as defined here becomes a property of the coding structure itself, that is, of the rules governing the mapping of the 4^*L*^ possible genotypes into phenotypes. Our formulation does not require the phenotype to be discrete and applies equally to continuous traits such as TF-binding affinity. The key requirement for its application is the knowledge of the full genotype-phenotype map, which is rarely accessible. However, it is tractable for many TF-binding motifs, where the mapping from sequence to TF-binding affinity can be modeled at nucleotide resolution (Mascolo et al. 2026), and for the genetic code, where the codon-amino acid mapping is known. These cases therefore provide a unique opportunity to analyze robustness at the level of the information-encoding structure.

Our approach can be formally reconciled with neutral network theory in the three idealized cases used in our analytical derivations (Eq. 9), two using only *SI* (but distributed differently across positions) and a full-*MI* motif (**Figure 2**). In these idealized cases, we require TF-binding sequences to be equiprobable. Under this simplifying condition, the phenotype becomes effectively binary: in these motifs, four 2-bp sequences bind the TF with equal probability, and the remaining twelve sequences do not bind it at all. Due to this discretization of the phenotype, the classic notion of robustness based on neutral networks can be applied to all three motifs. The resulting phenotype loss probabilities are different from the information loss rates we propose in this work, which depend on *N* and are expressed in *bits per mutation*. Nonetheless, phenotype loss probabilities recapitulate the 1:2 and 2:3 ratios we analytically derived for the asymptotic information loss rates (**Figure 7**). Moreover, this binarized limit case provides an intuitive illustration of why fully *SI*-encoded motifs can differ in robustness: robustness depends not only on the percent *SI*, but also on how *SI* is distributed across positions.

**Figure 7.**
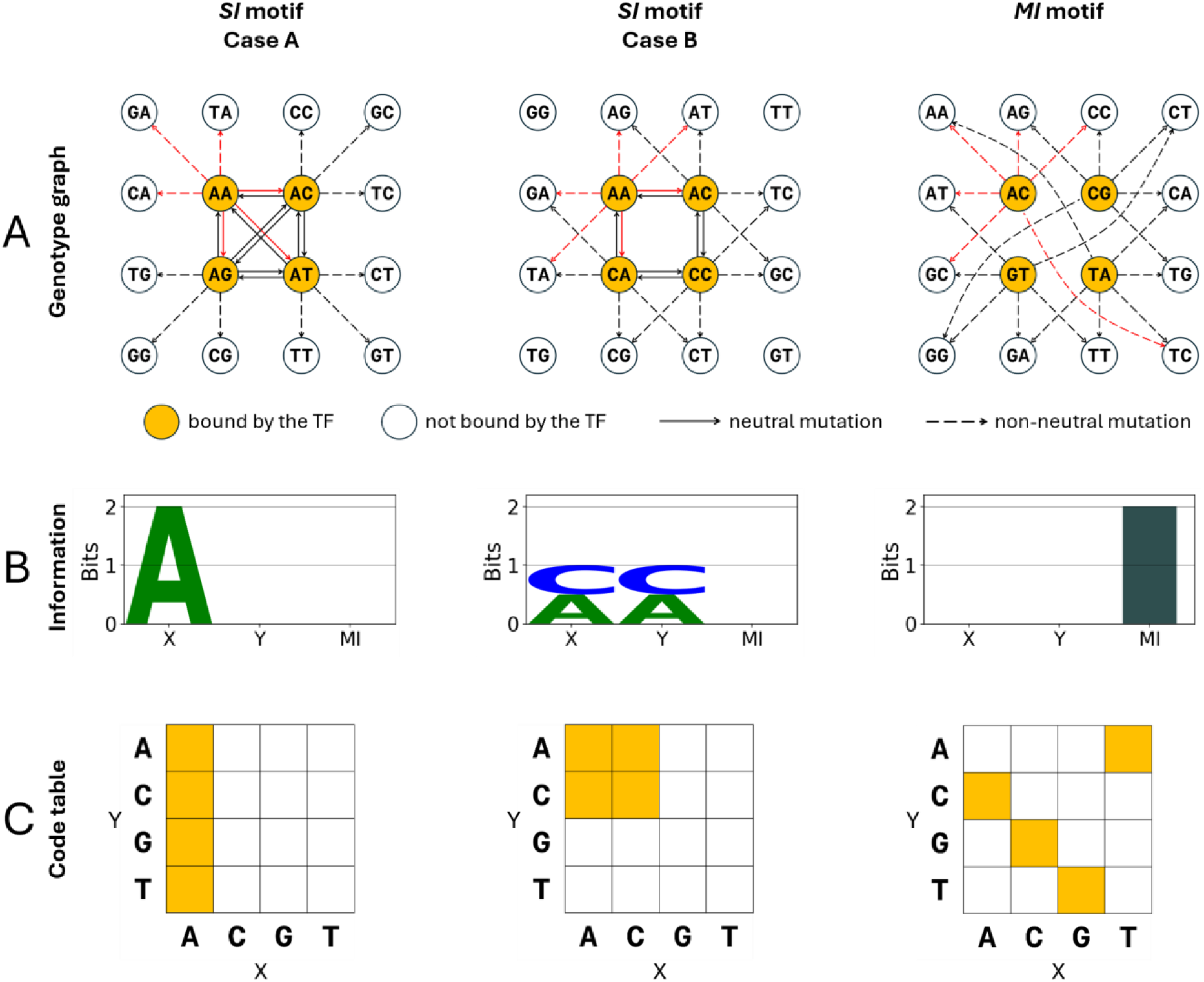
Comparison of three alternative descriptions of sequence motifs encoding a binary phenotype, applied to the three idealized motifs used in our analytical derivations, which represent three ways of encoding two bits at two positions (**Figure 2**). **(A)** Genotype graphs: the sequences composing the motif (i.e., bound by the TF) are represented with filled (yellow) circles, and can be mutated in six possible ways (three possible nucleotide substitutions at each position). Each mutation of a binding sequence can turn it into another binding sequence (solid arrows) or into a non-binding sequence (dashed arrows). For *SI* Case A (left), there are 12 deleterious and 12 neutral mutations. Thus, the fraction of deleterious mutations is 12/24=1/2. For *SI* Case B (middle), the fraction of deleterious mutations goes up to 16/24=2/3, while for the *MI* motif (right), all the 24 mutations are deleterious. For one chosen genotype in each graph, mutations are represented with red arrows to aid counting. **(B)** Sequence logos representing the information content of the motifs. The conventional representation of *SI* for the two positions (*X* and *Y*) is followed by an extra dark bar quantifying the amount of *MI* between X and Y (which is normally not included in sequence logos, since correlations are neglected). **(C)** Code tables where each sequence position (in this case, *X* and *Y*) is an axis, each with four possible entries. For the genetic code (L=3), the representation would be a 3D table (as in the ‘genetic cube’ in (Maeshiro and Kimura 1998)), but it is typically shown in 2D by partitioning each cell into four subcells (as in **Figure 6A**). In *SI*-based codes, encoded messages form (*L*-dimensional) rectangular subsets of the code table, up to permutations along each axis, but not in *MI*-based codes (see Appendix C).

More broadly, the perspective adopted in this work bridges bioinformatic descriptions of sequence motifs with evolutionary concepts such as neutral networks, showing that robust codes tend to be depleted in correlations among positions and enriched in *SI*. Somewhat counterintuitively, this hybrid approach also reveals that robustness can be further enhanced by concentrating *SI* in a few positions and hence reducing the mutational footprint of encoded information (Hledík et al. 2022). Notably, our results are in agreement with the high *SI* percentages and highly unequal distributions of *SI* observed in both TF-binding motifs and the genetic code (**Figure 6**C, left panel; (O’Neill and Erill 2016)).

### 3.2 Robust codes generate smooth, weakly epistatic landscapes

The distinction between *SI* and *MI* is not merely a statistical description of sequence motifs, but an information-theoretic formulation of the long-studied problem of epistasis in molecular evolution. Indeed, epistatic interactions require *MI*-based codes, since *SI* entails purely additive (non-epistatic) landscapes (see Appendix D). Previous work (Strelioff et al. 2010) demonstrated that in epistatic landscapes, epistatically interacting loci evolve *MI*, whereas in the absence of epistasis, sequences evolve *SI*-based motifs. But what happens if one allows molecular recognizers to coevolve with their target DNA sequences, so that the degree of epistasis is itself evolvable? Our results show that indirect selection for mutational robustness leads to *SI*-enriched (hence only weakly epistatic) sequence motifs, even in the absence of direct selection for *SI*, and despite a representational bias toward *MI* (**Figure 4**). While the specific equilibrium values depend on many system-specific evolutionary and population-genetic parameters (such as mutation rates, population size and the selection coefficients), we show that a preference for *SI* spontaneously evolves when the phenotype must be preserved in the face of mutations (**Figure 4**B).

This finding has implications beyond mutational robustness. Indeed, *MI*-based codes imply nonlinearities in the phenotype landscape (i.e. epistasis), including *reciprocal sign epistasis*, which is the type of interaction that can generate local optima and hinder navigability (Poelwijk et al. 2011; Saona et al. 2022) (**Figure *5***). In contrast, by avoiding epistasis, purely *SI*-based codes entail the smoothest possible landscape (a linear, or additive, landscape). In such landscapes, there is only one optimum, phenotypic and genotypic distances are strongly correlated, and beneficial mutations can be acquired in any order, thus maximizing the number of accessible paths to adaptation (**Figure 5**). Thus, the mutational robustness conferred by *SI*, to fit populations during periods of *stabilizing* selection, does not come at the expense of evolvability, which is instead crucial when the population undergoes *directional* selection (e.g., right after an environmental change). This is consistent with substantial evidence that, in some cases, robustness can facilitate adaptation rather than impede it (Masel and Trotter 2010; Draghi et al. 2010; Wagner 2007; Aldana et al. 2007).

The best example of a biological code that provides robustness and evolvability is the genetic code: it minimizes the effect of point mutations and mistranslations while facilitating adaptive exploration of protein sequence space (Maeshiro and Kimura 1998; Schwersensky et al. 2020; Rozhoňová et al. 2024). In a classic study (Maeshiro and Kimura 1998), Maeshiro and Kimura noted that the genetic code follows a specific encoding principle: synonymous codons are organized into 3D boxes within the “genetic cube” (i.e., the code viewed as a 3D table with the axes corresponding to the three codon positions). The only exceptions to this rule are the Leu, Arg, Ser and Stop codons. While their original formulation is difficult to parse and specific to the genetic code (*L*=3, 21 messages), our work provides a simple and general interpretation. In pure-*SI* motifs of length *L*, codewords encoding the same message necessarily form *L*-dimensional hyper-rectangles within the code table (see **Figure 7**C and Appendix C). The geometric organization identified by Maeshiro and Kimura essentially corresponds to *SI* encoding in the special case of the genetic code (*L*=3, 21 messages). Our work thus reveals that the genetic code closely follows a rather simple encoding strategy, the same as our most robust idealized motif (“*SI* motif, Case A”): encoding information solely as *SI* and packing it in a minimal set of highly conserved positions, making nucleotide contributions *independent* and *unequal*. The only four messages that depart from this strategy and involve correlations among positions are precisely the four exceptions identified by Maeshiro and Kimura (see Fig. **Figure 6**C, **Figure S10**).

### 3.3 Additivity as an evolved feature of biological codes

Classical models of TF-DNA interactions assume that TF-binding site positions provide independent contributions to the overall binding affinity. The scientific community has also explored higher-order models that incorporate correlations between positions, but they do not generally improve TF-binding site predictions substantially (Osada et al. 2004; Kulakovskiy et al. 2013; Omidi et al. 2017), and thus the PSFM remains the model of choice (Stormo 2013). This indicates that TF-binding motifs do not encode a substantial amount of information via correlations between positions, which raises the question: why make so little use of a valid source of information?

One possible explanation is that it would be difficult for proteins to recognize inter-positional correlations. However, observed correlations, especially between adjacent bases, are often implied by sequence-dependent structural features of DNA, such as local curvature or bendability, that proteins can readily recognize (Rohs et al. 2009; Velmurugu et al. 2018). Moreover, high-throughput experimental methods have been used to estimate global TF-DNA affinity landscapes with increasing accuracy and coverage (Maerkl and Quake 2007; Zhao et al. 2009; Aguilar-Rodríguez et al. 2017; Le et al. 2018; Westmann et al. 2024). These studies have shown that contributions to site occupancy are predominantly additive for high-affinity sites, whereas they consistently report non-additive contributions for low-affinity sequences. These observations support our interpretation that additivity might be largely an evolved feature of robust encodings, rather than a biochemical constraint on TF-DNA interactions. Indeed, after analyzing the standard genetic code - the best-known genotype-phenotype map, where any alternative tRNA-amino acid assignment would be biochemically possible - we have found the same qualitative trends observed for TF-binding motifs.

These results point to a general evolutionary pressure favoring independence among sequence elements. Consistent with this view, growing evidence suggests that, as with TF-binding motifs, the genotype-phenotype-fitness maps of protein-coding sequences are also more linear than previously assumed. After accounting for nonlinear transformations of the measured phenotype (sometimes referred to as non-specific or global epistasis), additive maps can explain most of the observed phenotypic variance, with the remaining unexplained variance being mostly due to pairwise specific epistasis, rather than higher-order (Park et al. 2024; Otwinowski et al. 2018; Grah et al. 2025; Weinreich et al. 2018; Carneiro and Hartl 2010; Metzger et al. 2024). Therefore, our framework rationalizes several recurring features of biological codes: nearly additive phenotypic effects, information-dense loci, and the coexistence of mutational robustness and evolvability. Importantly, this work recasts in information-theoretic terms several convergent insights that have independently emerged across traditionally distinct research communities, including bioinformatic approaches to sequence motifs and evolutionary genetic concepts such as robustness, evolvability, and epistasis. In doing so, it proposes a common framework for studying quantitatively how information encodings evolve in nucleic acid sequences.

## 4. Methods

### 4.1 Motif information characterization

For a TF-binding motif of length *L*, we compute the *single-locus information* as:

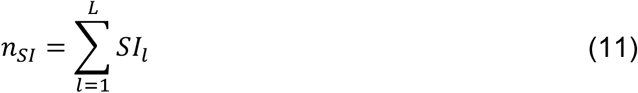

where *SI*_*l*_ represents the *single-locus information* at position *l*.

We compute the motif *mutual information* between adjacent positions as:

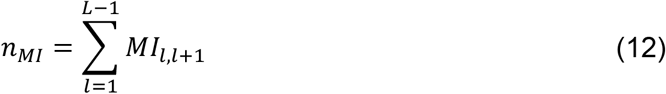

where *MI*_*l,l*+1_represents the Mutual Information between motif positions *l* and *l* + 1.

When we model TF binding preferences, we restrict the possible inter-positional interactions to pairwise interactions between adjacent positions (see Section 4.3 and Figure S10). This is done because contributions from higher-order correlations among positions, or non-adjacent pairwise interactions, are typically assumed to be very small. In this context, the motif’s total information available to the TF is simply:

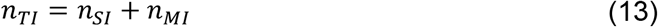

However, the true total information accounts for the total correlation *C* within all positions of the sequence motif (Eq. 7). This exact partition of *TI* into *SI* and *C* is used in the analysis of the genetic code (**Figure *6*** and **Figure S*10***), where the restriction to pairwise interactions is not appropriate, nor necessary. In this case, *n*_*TI*_ is calculated based on Eq. 7, rather than Eq. 13.

Finally, the *percent SI* is simply the percentage of information encoded via *SI*, that is, 100 · *n*_*SI*_/*n*_*TI*_.

### 4.2 Motif sampling

We apply two different algorithms to sample motifs with a required information content. When sampling from the full space of possible encodings (the general case), we apply an evolutionary algorithm (see Section 4.2.1). We also use this algorithm to generate motifs with specific values of *SI* and *MI*. When instead we work under the restricting assumption that selected (target) sequences are equiprobable, we can exactly enumerate all motifs and directly draw uniform samples (see Section 4.2.2).

#### 4.2.1 Evolutionary algorithm for sampling generic motifs

For the general case, we employ an evolutionary algorithm to sample TF-binding motifs encoding a specific amount of *SI* and/or *MI*. Motifs are explicitly represented as a collection of *n*_*bs*_ TF-binding sites of a given length (*l*_*bs*_). The evolutionary algorithm is initialized with a population of randomly initialized motifs and a requested number of bits of *SI* (*r*_*SI*_) and/or *MI* (*r*_*MI*_). The algorithm then iterates a replication-mutation-selection process, using the following error function as the objective function to be minimized:

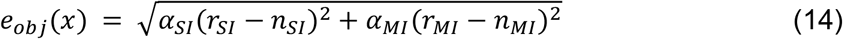

where *α*_*SI*_ and *α*_*MI*_ are control (0,1) values determining whether *SI* (*r*_*PI*_) and/or *MI* (*r*_*MI*_) are included as targets in the error function to be optimized.

At each iteration, each individual motif is first duplicated, and mutations are applied to its copy by randomly replacing nucleotides at each site position with a probability of 0.01 per position. The algorithm then compares the *error e*_*obj*_(*x*) of the parent and offspring motifs in terms of their deviation from the desired number of bits of *SI* (*r*_*SI*_) and/or *MI* (*r*_*MI*_), and the motif with the lower error proceeds to the next generation.

We principally employ *α*_*SI*_ = 1, *α*_*MI*_ = 0 and *α*_*SI*_ = 0, *α*_*MI*_ = 1 to generate *SI-* or *MI*-based motifs, respectively. However, we use *α*_*SI*_ = 1, *α*_*MI*_ = 1 to explicitly control for the fraction of both *SI* and *MI* in specific cases (**Figure S2**).

#### 4.2.2 Exact sampling of motifs with equiprobable target sequences

Under the assumption that target sequences are equiprobable, we can exactly enumerate and sample motifs that have a required total information content (*TI*). This restrictive assumption, which was used in defining the three idealized motifs in Section 2.1, effectively induces a binary phenotype in which *n*_*selected*_ *L*-mers are bound with equal probability, while all the others are unbound. Under this assumption, *TI* = *log* (4^*L*^/_*selected*_). Motifs are then obtained by uniformly sampling over the 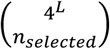 possible subsets of recognized sequences. This approach was used to estimate the probability distributions in Fig. 4C, S7 and S8.

### 4.3 Mutational robustness analysis

#### Information Loss Analysis

To analyze information loss in motifs, we sample two sets of 100 motifs to encode information as either *SI* or *MI*, or a specific mixture of both in **Figure S2**. We then iteratively apply mutations, randomly substituting in each motif a single base of the constituent *n*_*bs*_ TF-binding sites at each iteration, and monitoring the average *SI* and *MI* in each motif set as iterations progress.

#### Network Integrity Analysis

To analyze network integrity, we sample two sets of 100 motifs to encode information as either *SI* or *MI*. For each sampled motif, we generate a random genome sequence of length *l*_*G,int*_, and the *n*_*bs*_ TF-binding sites that define the motif are then embedded into this sequence. The embedded TF-binding sites are used to instantiate a dinucleotide PWM (diPWM) model (Kulakovskiy et al. 2013), which accounts for dependencies among adjacent nucleotides, and the diPWM model is used to calculate TF-binding affinity scores along the artificial genome sequence using a conventional sliding window approach. The embedded TF-binding sites are the true target sites, while all the other genomic positions are defined as non-target sites. The ability of the model to distinguish target sites from the rest of the genome sequence is captured by the area under the precision-recall curve (AUPRC). We iteratively apply mutations, randomly substituting a single base of the genome at each iteration, and monitoring the average AUPRC for each motif set as iterations progress.

#### Evolutionary simulations

To explore the mutational robustness of TRNs, we simulate the coevolution of TF-binding motifs and their target sites within an artificial genome. Each organism in this co-evolutionary algorithm is defined by a genome that is represented as a DNA sequence. The locations for target sites are pre-designated, and the TF-binding model is stored in a pre-specified ‘gene’ region of the genome, where the diPWM model is represented as a set of TF-binding sites arranged column-wise **Figure S*11***. The gene region is read at each generation to instantiate a diPWM model, which is then used to scan the genome and create a vector of binding scores representing the binding affinity for each position in the genome. The positions designated as TF-binding sites are the true target sites, while all the other genomic positions are defined as non-target sites. The fitness of each organism is then defined by its classification performance, as captured by the AUPRC:

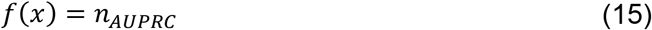

where *n*_*AUPRC*_ is the organism’s AUPRC.

The algorithm evolves a population of organisms under fitness function *f*(*x*) following, at each iteration, a three-step process (Tenaillon et al. 1999). During *reproduction, n*_*offspring*_ copies of each organism are made based on the organism’s fitness *f*. In particular, *n*_*offspring*_ is drawn from a Poisson distribution with expectation defined by the relative fitness of the individual:

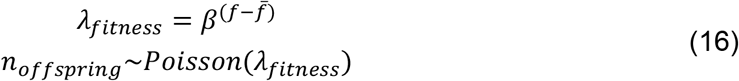

where, 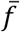 is the average fitness of the population and *β* modulates the strength of selection.

During *mutation*, parents and offspring are all mutated by randomly applying one single-nucleotide substitution in the genome of each organism. Lastly, during *sampling*, drift is modeled by randomly sampling the population to maintain the population size constant.

#### Seeding of evolutionary simulations

To seed evolutionary simulations at specific points in motif space (**Figure 4AB, Figure 5**), we use the same coevolutionary framework, but we introduce an error term, *e*_*seed*_(*x*), to the fitness function in order to enable selection for a specific proportion of single-locus information (*SI*) as part of the motif total information (*TI*), as well as for specific values of AUPRC:

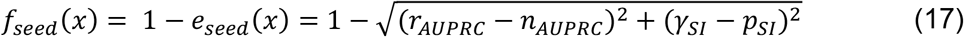

where *n*_*AUPRC*_ is the organism’s AUPRC, *r*_*AUPRC*_ is the target AUPRC, 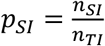 is the proportion of the organism’s *TI* (*n*_*TI*_) encoded through *SI* (*n*_*SI*_) and *γ*_*SI*_ is the requested proportion of total information to be encoded through *SI*.

#### TRN competition

To evaluate the mutational robustness of *SI*-encoded networks versus *MI*-encoded networks, we first generate seed organisms of each class by running the coevolutionary simulation algorithm with the modified fitness function *f*_*seed*_(*x*). For both classes, we select for perfect network functionality (*r*_*AUPRC*_ = 1), and we apply *γ*_*SI*_ = 0.70 for seed organisms with *SI*-encoded networks, and *γ*_*SI*_ = 0.30 for seed organisms with *MI*-encoded networks. In each simulation, we select the first organism to reach AUPRC=1, and selected organisms of each class (25 *SI* organisms and 25 *MI* organisms) are then randomly paired with replacement to create 100 *MI*/*SI*-based organism pairs. For each pair, we use the selected organisms to create a clonal population of 2,000 seed organisms (1,000 for each encoding strategy). This seed population is then used as the starting point for an evolutionary simulation using fitness function *f*(*x*), and we track the average AUPRC of each strategy as the simulation progresses.

#### Vector Field analysis

To analyze the mutational and selective constraints of the coevolutionary simulation, we divide the phenotype space of TRNs into a grid of 25 cells spanning all combinations of 5 bins in terms of AUPRC and %*SI*. For each cell, we generate seed organisms by running coevolutionary simulations with a randomly initialized population of 2,000 organisms, adjusting *γ*_*SI*_ and *r*_*AUPRC*_ in the *f*_*seed*_(*x*) function to target the center position of the cell. These simulations are run until one organism of the population reaches a Euclidean distance from the cell center of at most 0.01 (*f*_*seed*_(*x*) ≥ 0.99), at which point we select the organism as a seed organism and we clone it 2000 times to create a seed population for the cell. We use this seed population as the starting point for an evolutionary simulation using fitness function *f*(*x*). We stop the simulation when the population’s centroid reaches the cell’s boundary. This process is repeated ten times per cell, yielding 10 independent replicas of the evolutionary experiment in each cell and, for each cell, we calculate the average displacement vector, which is shown in **Figure *4*B**.

We generate the evolutionary steady state by rerunning the coevolutionary simulation for each of the seed populations in the (70% *SI*, 0.9 AUPRC) cell. We allow the simulation to proceed without stopping regardless of whether cell boundaries are crossed. After 27,000 generations, we report the achieved steady state as the average of the ten population centroids (visualized as the black star in **Figure *4*B**). We generate the mutational steady state through a similar procedure. We run the coevolutionary simulation for each seed population in the (30% *SI*, 0.1 AUPRC) cell without selective pressure for 10,000 generations, and we report the achieved steady state as the average of the ten population centroids (visualized as the black cross in **Figure *4***B).

### 4.4 Affinity landscape analysis

In our study of the genotype-phenotype map, we associate each genotype *x*_*i*_ with a phenotype *f*(*x*_*i*_), namely the TF-DNA binding affinity for the DNA sequence *x*_*i*_. The affinity is scored with the standard log-likelihood ratio approach (Hertz and Stormo 1999). Importantly, the score is assigned using the true joint probability distribution of the *L*-mers: *f*(*x*_*i*_) = *log*_2_(*p*(*x*_*i*_)/*p*_0_(*x*_*i*_)), where *p*(*x*_*i*_) is the probability of a selected site being *L*-mer *x*_*i*_, while the genomic background probability for that *L*-mer is *p*_0_(*x*_*i*_). Only in the special case of pure *SI* motifs the joint distribution factorizes and this expression reduces exactly to the standard additive PSSM score, while for motifs that contain *MI* it correctly captures the epistatic structure that a PSSM approximation would erase. Genotype distances are simply defined as the Hamming distance (Hamming 1950), i.e., the number of positions where the sequences differ. Two DNA sequences are *adjacent* to each other - or *neighbors* - if they are at a Hamming distance of 1 from each other (i.e., one sequence can be turned into the other by a single nucleotide substitution).

#### FDC

The fitness-distance correlation (FDC) is a widely used landscape metric to study evolvability, developed to estimate the navigability of fitness landscapes (Sun et al. 2014; Zou et al. 2022). It measures the correlation between the fitness of a genotype and its distance (in genotype space) from the global optimum. Here we apply this metric to estimate the navigability of the affinity landscape, so *f*(*x*) is the TF binding affinity for sequence *x*, and not its fitness. If we denote as *d*(*x, o*) the (Hamming) distance of a genotype *x* from the global optimum *o* (the sequence with the highest affinity), the FDC is defined as

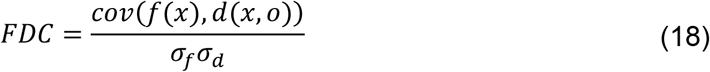

where *σ*_*f*_ and *σ*_*d*_ are the standard deviations of *f*(*x*) and *d*(*x, o*), respectively. Bigger negative correlation coefficients indicate more navigable landscapes.

#### GDPDC

The genotype distance-phenotype distance correlation (GDPDC) is analogous (aside from a linear transformation) to a convolution of the FDC over all the possible pairs of genotypes (i.e., it is not restricted to pairs where one of the two elements is the global optimum). Indeed, for all the possible pairs of genotypes (*x*_*i*_, *x*_*j*_), it measures the correlation between their distance in genotype space *d*(*x*_*i*_, *x*_*j*_) and their distance in phenotype space *v* = |*f*(*x*_*i*_) ™ *f*(*x*_*j*_)|. Thus,

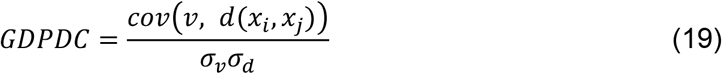

where *σ*_*v*_ and *σ*_*d*_ are the standard deviations of *v* and *d*(*x*_*i*_, *x*_*j*_), respectively. Higher correlation coefficients indicate more navigable landscapes (Greenbury et al. 2016).

#### Number of optima

A set of genotypes *O* that forms a *connected component* in the genotype graph (the graph that links neighbor genotypes) is an optimum if and only if

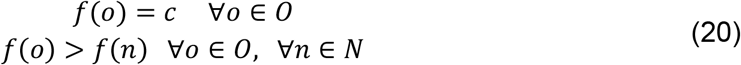

where *c* is a constant, and *N* is the set of neighbors of the elements of *O* that do not belong to *O* (Stadler 2002). The set *O* can contain a single element if that single genotype has a higher phenotype value than all its neighbors. When calculating the FDC, if the global optimum is a set of more than one genotype, *d*(*x, o*) is defined as the distance from the closest genotype that belongs to the optimum *O*. The number of optima is simply the total count of optima in the whole genotype-phenotype map.

## Code availability

The code for all the simulations and analyses presented in this study, including the generation of figures, is freely available at https://github.com/ErillLab/ITTF.

## Acknowledgments

This research was supported in part by the Spanish Ministry of Science, Innovation and Universities (MICIU) under grants PID2023-152240OB-C21 and PID2024-156292OB-I00. The computational studies reported in this work have been performed using the UMBC High Performance Computing Facility (HPCF). The facility is supported by the U.S. National Science Foundation through the MRI program (CNS-0821258, CNS-1228778, OAC-1726023 and CNS-1920079) and the SCREMS program (DMS-0821311), with additional support from the University of Maryland Baltimore County (UMBC). Emmanuel Mekasha and Elia Mascolo were funded by the Merck Data Science for Observational Research Program through the UMBC College of Natural and Mathematical Sciences Academic Fellowship Program. Elia Mascolo is a recipient of an APART-USA Fellowship of the Austrian Academy of Sciences at the Institute of Science and Technology Austria (ISTA).

The authors thank Gašper Tkačik and Aleksa Spasić for helpful feedback on a draft version of the manuscript.

## Author contributions

### Conceptualization and Supervision

Ivan Erill; Elia Mascolo.

### Methodology, Software, Investigation, Formal analysis, Visualization

Emmanuel Mekasha; Elia Mascolo.

### Writing - Original Draft

Elia Mascolo; Ivan Erill; Emmanuel Mekasha.

### Writing - Review & Editing

Ivan Erill; Elia Mascolo; Emmanuel Mekasha.

### Project administration

Ivan Erill.

## Appendix A. Information-theoretic framework

### Appendix A1. Estimating total information

The most general definition of biological information is the Kullback-Leibler (KL) divergence between the observed genotype distribution *P*(*σ*) and the null genotype distribution *Q*(*σ*), which is the expected distribution in the absence of selection. Thus, the total information (*TI*) at a given *L*-bp genomic location is

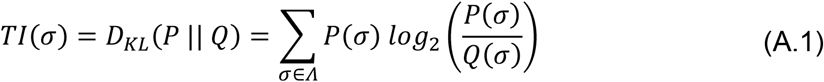

where *Λ* is the set of all the possible *L*-mers.

A common approximation (Strelioff et al. 2010) assumes a uniform null distribution. Under this assumption, *Q*(*σ*) becomes simply a constant, equal to 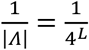, and the *TI* thus reduces to

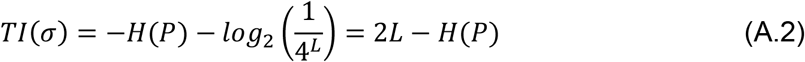

where *H*(*P*) is the entropy of the observed genotype distribution. Note that, when the null distribution is uniform, its entropy is *H*(*Q*) = 2*L* bits, so that the information is simply a difference between two entropies.

This metric is often applied to calculate the information of a genomic locus. In this case, *P*(*σ*) is the distribution for the genomic sequence *σ* over all the observed genomes in a population. However, the same metric is routinely used to calculate the information carried by the sequences targeted by a molecular recognizer within one single genome. In that context, *P*(*σ*) is the distribution over the recognized sequences in the genome. Then Eq. A.2 can be re-written as

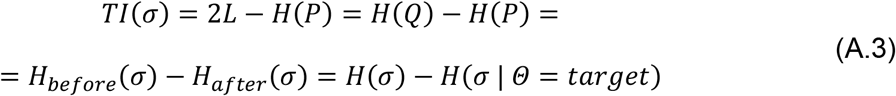

where *H*_*before*_(*σ*) is the *a priori* uncertainty on the sequence, while *H*_*after*_(*σ*) is the *a posteriori* uncertainty, given that the sequence is one of the recognized targets (i.e., is drawn from distribution *P* rather than *Q*) (T. D. Schneider et al. 1986). Following Shannon’s definition, Eq. A.3 measures the information gained on *σ* after receiving the message *Θ* = *target* (i.e., how much the uncertainty on the sequence decreases once we know it is one of the targets) (Shannon 1948). Therefore, under a uniform null distribution *Q ~ U*, the *TI* of a sequence motif is simply a difference in entropy.

#### Appendix A.2. Decomposing total information into single-locus information and correlations between loci

Following a standard approach (Eq. A.3; (Strelioff et al. 2010)), the total information content of an *L*-bp sequence motif (*σ*) can be expressed as *TI*(*σ*) = *H*_*before*_(σ) ™ *H*_*after*_(σ). In the prior distribution, all nucleotide positions (*σ*_*i*_) are assumed to be independently uniformly distributed, so that

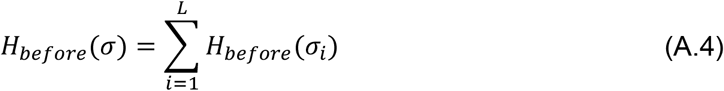

However, in the observed joint distribution (*σ*_1_, *σ*_2_, …, *σ*_*L*_), there can be statistical dependencies among the *σ*_*i*_ variables. These correlations can be measured using the *total correlation*, which is a generalization of the mutual information to more than two variables [11]. Indeed, analogously to how *MI* can be expressed as the KL divergence from statistical independence, *MI*(*X*; *Y*) = *D*_*KL*_(*p*(*X, Y*) || *p*_*X*_ · *p*_*Y*_), the total correlation between *L* random variables (*σ*_1_ to *σ*_*L*_) is

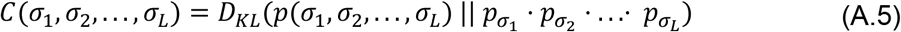

which reduces to 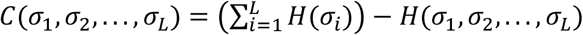. Therefore,

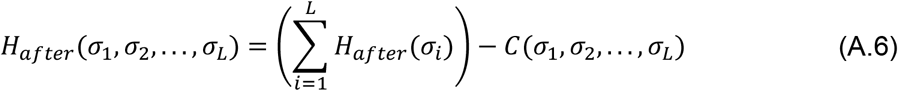

We can now rewrite the total information as follows:

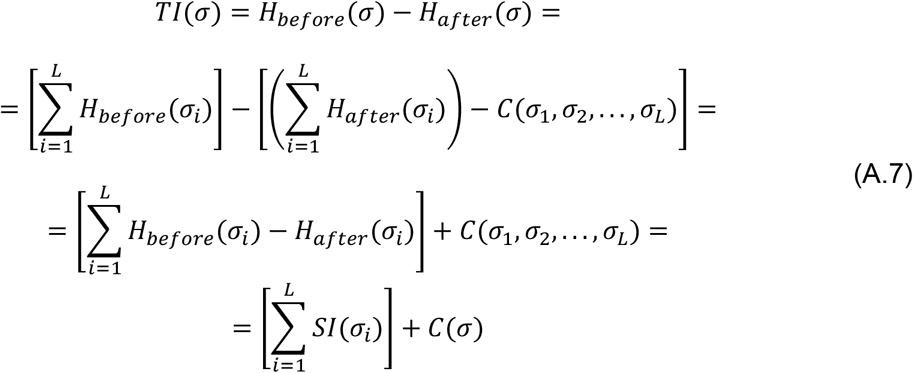

Therefore, any code storing information in *L*-bp DNA sequences will necessarily encode a fraction *p* of the total information through single-locus information (*SI*) and the remaining fraction, 1 ™ *p*, through the correlations among the *L* positions, with 0 ≤ *p* ≤ 1.

In the special case of *L*=2, the decomposition simply reduces to the sum of *SI* and *MI*:

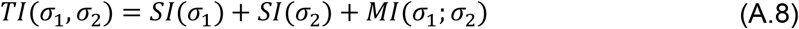

## Appendix B. Asymptotic information loss due to mutation

### Appendix B.1. Approach

Mutations affect information content by altering the probability distribution over nucleotide sequences underlying a sequence motif. *SI* and *MI* are smooth functionals of the underlying probability distribution, so we can analytically characterize their sensitivity to perturbations of the probability distribution. However, the sensitivity also depends on several properties of the probability distribution, such as the total information content and how it is distributed across the marginal probability distributions (i.e., across each position of the sequence motif). Thus, to perform a meaningful comparison, we first studied *SI* and *MI* motifs under the following restrictions: (1) all the compared motifs have the same information content *TI*/*L* = 1 *bit per position* (approximately what is found in actual TF motifs); (2) the target sites recognized by the TF are all equally represented (so they are equiprobable).

Given that MI requires two distinct positions to be correlated, the shortest motif length to serve as model for the *SI*-*MI* comparison is 2 bp. Therefore, we compared *SI* and *MI* for simple motifs where *L*=2. Due to restriction (2), the total information is 2 bits in each motif. We require the 2 bits to be encoded in the two motif positions either purely via *MI* or purely via *SI*.

#### Pure MI motif

For *MI*, encoding 2 bits under the above-mentioned conditions requires selecting 4 out of the 4^L^ = 16 possible DNA dimers (*TI* = *log*_2_(16/4) = 2 bits), in such a way that all four nucleotides are represented at each position, yielding

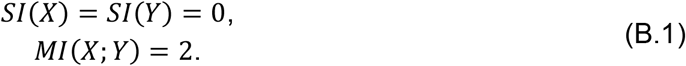

An example of such a motif is shown in **Figure 2C**.

##### Pure SI motif

For *SI*, the information can be distributed asymmetrically over the first position (*X*) and the second position (*Y*), as long as *SI*(*X*) + *SI*(*Y*) = 2 and *MI*(*X*; *Y*) = 0. To account for this, we studied the two extreme cases:

##### Case A

Information is distributed as unequally as possible over the motif, i.e., entirely carried by one position. That position will carry all 2 bits by being fully conserved (selecting one nucleotide out of four), while the other position will carry no information by having all four nucleotides equally represented. Without loss of generality, we can choose the conserved position to be the first one (*X*), leading to

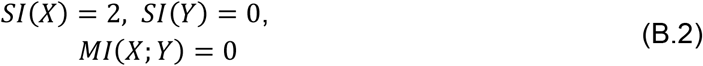

An example of such a motif is shown in **Figure 2A**.

##### Case B

Information is distributed as equally as possible over the motif, i.e., 1 bit in each position. This implies selecting 2 out of the 4 possible nucleotides at each position, while avoiding correlations, i.e., by requiring *P*_*XY*_ (*x, y*) = *P*_*X*_(*x*) · *P*_*Y*_(*y*), leading to:

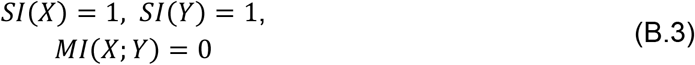

An example of such a motif is shown in **Figure 2B**.

#### Expected information loss due to mutation

Let *P* denote the motif distribution before mutation and *P*^′^ after mutation. We study the smallest mutational change possible, namely the random replacement of just one nucleotide (chosen uniformly at random) of one of the *N* aligned sites composing the TF motif (chosen uniformly at random) by one of the other three nucleotides, each with probability 1/3. This change corresponds to a single nucleotide substitution. The information loss is measured by Δ*I* = *I*(*P*′) ™ *I*(*P*). To estimate the robustness of the information-encoding strategies, we study their expected loss E[Δ*I*]. The expected loss due to mutation is a function of *N*, because a single nucleotide substitution will induce a perturbation *ε* of the probability distribution of order *O*(1/*N*). Taking the limit for *N* → ∞ is informative, as it provides the derivative of the information functional with respect to mutational perturbation:

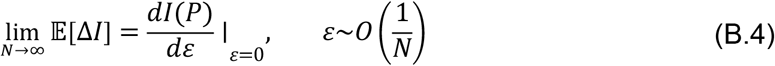

This asymptotic information loss serves as a local estimator of the sensitivity of information-encoding strategies of interest to mutation. Moreover, it is an increasingly good approximation of the expected loss E[Δ*I*] as *N* grows.

Therefore, we proceed to analyze the information loss rates for the three motifs described above, namely the *MI*-based motif and the two extreme cases (case A and case B) for *SI*, which are named, respectively:

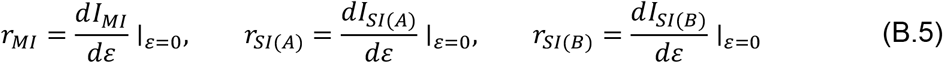

#### Appendix B.2. Information loss calculations

##### Appendix B.2.1 Loss rate for *SI* motif, Case A

For Case A, two mutational events are possible: the mutation can hit the position carrying the fully conserved nucleotide; the mutation can hit the other position where the distribution over the four nucleotides is uniform. We will assume, without loss of generality, that the fully conserved position is position *X*, and that the uniformly distributed one is position *Y*.

##### Mutation occurs at position *X*

A mutation will decrease by one the count for the conserved nucleotide, decreasing its frequency to 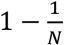. The nucleotide is mutated into a new one that now appears with frequency 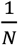 Therefore, the entropy after mutation is

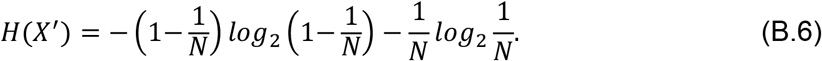

Using Taylor’s expansion for log_2_(1 ™ *ε*) on the first term we find

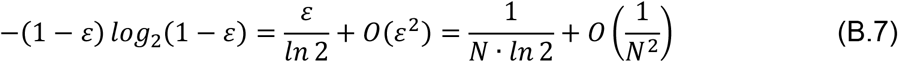

Therefore:

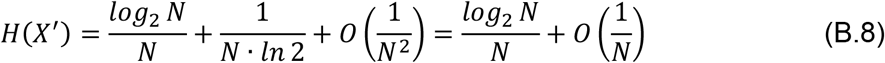

Since *H*(*X*) = 0,

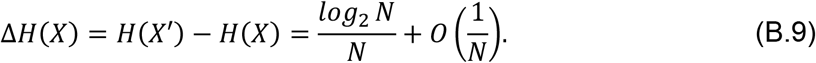

The information loss is

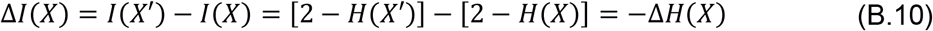

Therefore,

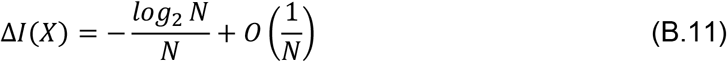

##### Mutation occurs at position *Y*

Position Y is uniformly distributed. A point mutation redistributing probability mass among an already uniform distribution leaves the entropy unchanged to leading order.

This can be proven by considering the entropy after mutation:

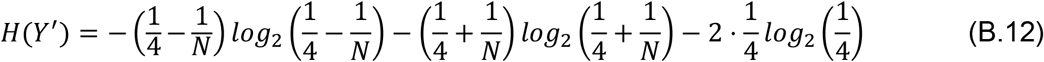

We use the Taylor expansion for small *ε*:

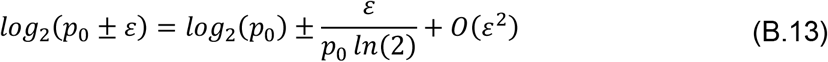

Applying it to the logarithms in Eq. B.12, where *ε* = 1/*N*, and *p*_0_ = 1/4, they become:

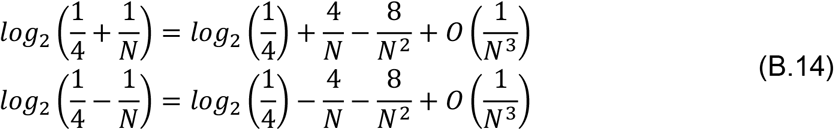

By plugging them into Eq. B.12 we obtain

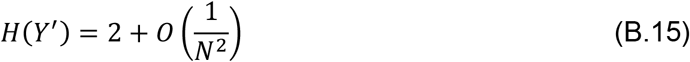

Since *H*(*Y*) = 2 bits, the information loss is

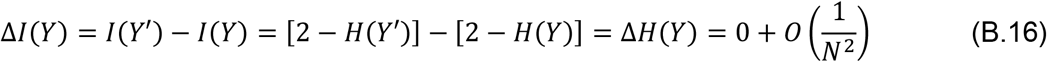

##### Expected value

The two possible mutational events (i.e., mutation hits the conserved position *X* or the non-conserved position *Y*), whose effects are measured by Eq. B.11 and B.16, can both occur with probability 1/2. Therefore, the expected information loss for *SI*-based motifs structured according to *Case A* is

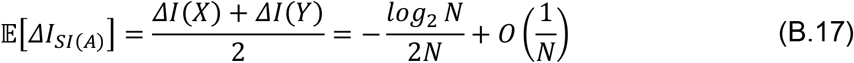

and the asymptotic information loss is

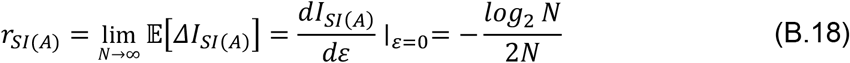

##### Appendix B.2.2 Loss rate for *SI* motif, Case B

For Case B, positions *X* and *Y* are identically distributed. Therefore, without loss of generality, we can just study the loss when a mutation hits position *X*.

At position *X*, two letters are already represented (with equal probability). A mutation is assumed to turn a nucleotide into one of the other three nucleotides with equal probability (1/3 each). There are two possibilities to consider: the mutation can turn a nucleotide into the other already represented letter (with probability 1/3); the mutation can turn a nucleotide into one of the two letters that were not already represented (with probability 2/3). As an illustration, the reader can consider the motif shown in **Figure 2B**, and suppose that a mutation hits an “A” at position *X*: the effect on the probability distribution will depend on whether the “A” is turned into “C” (the probability is 1/3) or into one of the non-represented nucleotides, namely “G” or “T” (the probability is 2/3).

##### Mutation to an already represented nucleotide

Analogously to the situation analyzed in Eq. B.12-16, a mutation redistributing probability mass among two equally likely states is only quadratic in 1/*N*, leaving the entropy unchanged up to *O*(1/*N*^2^). This can be proven by considering the entropy after mutation:

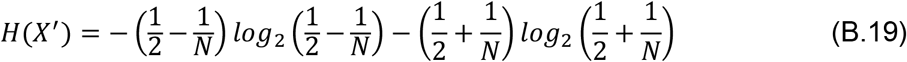

As done for the *SI* motif Case A, we can use Taylor expansions for the logarithms

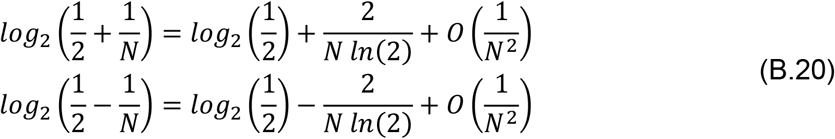

By plugging them into Eq. B.19 we obtain

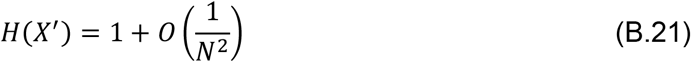

Since *H*(*X*) = 1, the information loss is

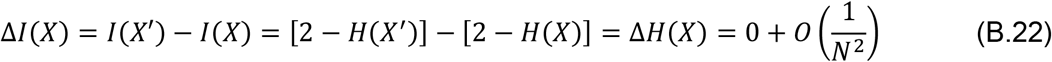

##### Mutation to a new nucleotide

When the mutation produces one of the two nucleotides that were not already represented the entropy becomes

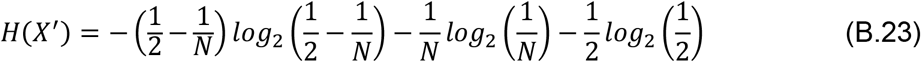

The first term can be rewritten, via Taylor expansion, as follows:

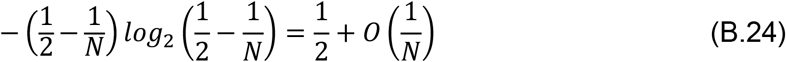

yielding

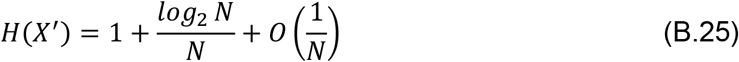

Since *H*(*X*) = 1, the information loss is

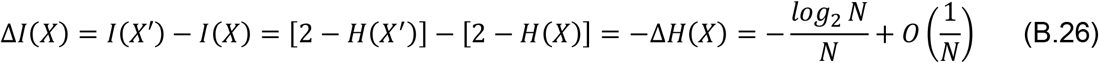

##### Expected value

The two possible mutational events (mutation to an already represented nucleotide or to a new nucleotide), whose effects are described by Eq. B.22 and B.26, can occur with probabilities 1/3 and 2/3, respectively. Therefore, the expected information loss for *SI*-based motifs structured according to *Case B* is

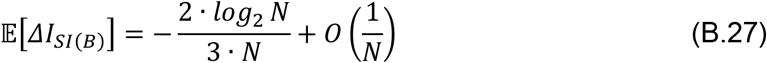

and the asymptotic information loss is

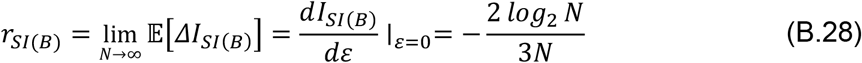

#### B.2.3 Loss rate for *MI* motif

The *MI*-based motif defined in Appendix B.1 encodes 2 bits by selecting (with equal probability of 1/4) four out of 16 possible DNA dimers (x,y). A mutation will decrease by one the count for one of the four represented dimers, decreasing its frequency from 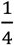 down to 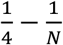. The dimer is turned into a different one: the new “mutated” dimer. By construction, the motif exhibits maximal correlation between the positions X and Y. In other words, knowing the letter at position X fully specifies which of the four letters is present at position Y (thus providing 2 bits of mutual information), and vice versa. Therefore, for any given letter at one position, there is only one acceptable letter at the other position. This implies that a single nucleotide substitution will always generate a non-selected dimer, i.e., one among the 12 that were not represented (whose frequency therefore rises from 0 to 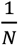). **Figure 2C** provides an example of a possible *MI*-based motif that encodes 2 bits, for illustration purposes. For three of the dimers already represented before mutation, the frequency remains 1/*N*. Overall, the entropy after mutation is

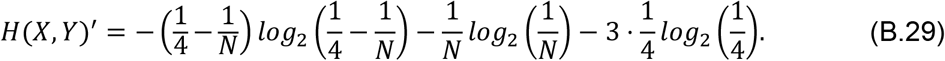

Therefore,

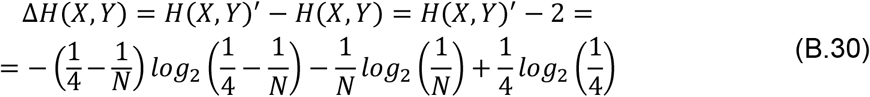

For the logarithm in the first term, we can use the Taylor expansion 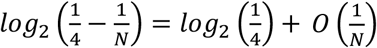, and plug it into Eq. B.29, obtaining

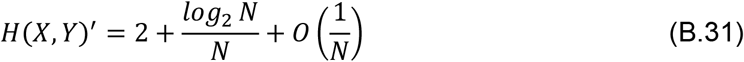

The entropy before the mutation is *H*(*X, Y*) = 2. Therefore,

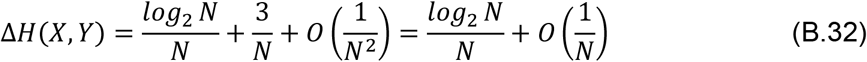

Since *MI*(*X*; *Y*) = *H*(*X*) + *H*(*Y*) ™ *H*(*X, Y*), we can rewrite Δ*I*_*MI*_ as

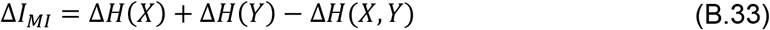

However, 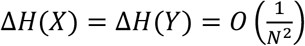 (because a mutation would redistribute probability mass between two equiprobable symbols of the uniform marginal distribution; see Eq. B.12-16). Therefore, the expected information loss for the *MI*-based motif is

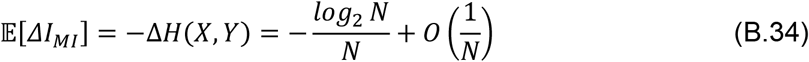

and the asymptotic information loss is

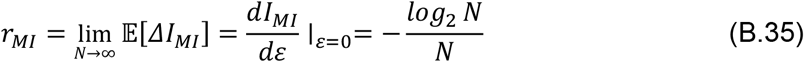

#### Appendix B.3. Comparison of loss rates

From Eq. B.18, B.28, B.35, we can see that the asymptotic loss rates for *MI*- and *SI*-based motifs are substantially different, and we find that

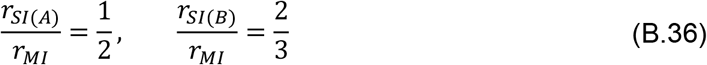

## Appendix C. *SI*-encoded messages represent rectangular subsets of the genotype table

There are 4^*L*^ possible *L*-mers. They can be organized into an *L*-dimensional *genotype table* with *L* axes, each with four possible entries (for the four possible nucleotides). Every cell in this table represents a distinct *L*-mer. Thus, the table can be represented as an *L*-dimensional (hyper)-rectangle, and the boundaries between rows and columns slice it in four parts along each dimension (see, as an example, the visualizations for the case *L*=2 in **Figure *7*C**, or see **Figure S*10*** for the case *L*=3).

When positions contribute independently of each other to molecular recognition (no correlations among positions), the probability distribution over the recognized codewords encoding a message follows a simple constraint: the probability that the recognized *L*-mer is sequence *B* simply factorizes as 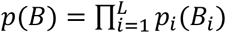, where *B*_*i*_ is the *i*-th element of *B*, and *p*_*i*_(*B*_*i*_) is the marginal probability of its occurrence at position *i* among recognized codewords encoding the message. If we now sort each axis of the table based on ascending or descending *p*_*i*_, one corner of the (hyper)-rectangle will contain a cell corresponding to an *L*-mer with the maximum probability, while the opposite corner the minimum probability, and probability monotonically increases/decreases in every direction.

When we also assume equiprobability among recognized codewords, the structure becomes even more constrained, because molecular recognition is effectively reduced to a binary yes-or-no phenotype. At each position *i*, only *N*_*i*_ nucleotide bases out of four are accepted (0 ≤ *N*_*i*_ ≤ 4), each with probability 1/*N*_*i*_, while the remaining ones are recognized with a probability of 0. Under this assumption, the recognized codewords are all equiprobable, with probability 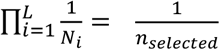, where 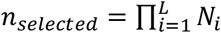 is the number of recognized *L*-mers. Due to the sorting along each axis, the marginal probability over each dimension *i* of the table will have *N*_*i*_ adjacent non-zero values (all with probability ^1^), followed or preceded by 4 – *N*_*i*_ zero values. Therefore, recognized *L*-mers form an *L*-dimensional rectangular subset of the (hyper)-rectangle with dimensions (*N*_*i*_ × *N*_*i*+1_ × …× *N*_*L*_) (shown in yellow in **Figure *7*C** *left* and *middle*). Deviations from this rectangular structure imply a violation of independence, meaning that 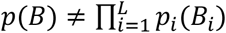, and thus the total correlation among positions is non-zero (**Figure *7*C** *right*; see also boxed motifs in **Figure S*10***).

## Appendix D. Pure-*SI* motifs imply additive (non-epistatic) landscapes

Two mutations are said to interact epistatically when the phenotypic effect of one mutation alone differs from its effect in the presence of the other mutation. In contrast, the absence of epistasis implies a purely additive interaction, such that

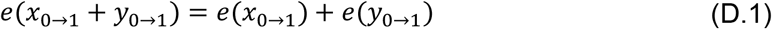

where 0→1 indicates a mutation from the initial nucleotide (marked with 0) to a new nucleotide (marked as 1), and the *e* function is the effect of mutations (a measurable phenotype). The various categories in which epistatic interactions can be classified represent different ways of departing from this equality.

When a phenotype is genetically encoded via *SI*, the DNA sequence positions have independent (purely additive) effects. Under independence, the joint distribution over all the *L*-mers is simply the product of the marginals at each position, and the information encoded in DNA becomes strictly additive (see Eq. 4 in the main text). In biological codes, sequence motifs are typically decoded by molecular recognizers that differentially bind the possible *L*-mers. This allows the uncertainty on any phenotype that relies on recognizer-DNA binding to be decreased by up to *TI*(*σ*) bits (see Eq. A.3), which is an upper bound on phenotypic information gain due to the *data processing inequality* theorem.

Epistatic interactions among sequence positions are expected to result in correlations (*MI*) (Strelioff et al. 2010). Instead, TF-binding sites are typically assumed to be approximately in a regime of independent effects from each nucleotide or with at most a few significant dinucleotide interactions (mostly from adjacent positions). This independent regime, captured by the conventional PSSM models, reduces the affinity score (based on the log-likelihood ratio; see **Figure *1***f) to a purely additive score. For example, if the *e* function in Eq. D.1 were to represent the classic PSSM score (**Figure *1***e-f), the equality would always hold true. Thus, when TF-DNA interactions are estimated using this classical bioinformatic model, the underlying assumption is that there are no significant epistatic interactions among the positions of the sequence motif. Instead, in the presence of epistatic interactions, selection stores genetic information also through *MI* (or higher-order correlations) among positions (Strelioff et al. 2010), breaking additivity (both sequence information contributions and binding energy contributions are no longer merely additive).

## Appendix E. The maximum number of optima for nucleotide dimers is four

Two optima in the landscape cannot be adjacent (only one mutation away from each other). Indeed, if they had both the maximum value, they would count as one (flat) optimum (see optimum definition in Methods, Section 4.4), while if they had different values, the lower one could not be an optimum, since it would have a neighbor with a higher value. Therefore, two optima always differ in at least two positions (for dimers, both positions must differ).

If four dimers are distinct optima, they must all have distinct nucleotides at position 1. With only four nucleotides available, this bounds the number of optima at four (position 2 gives the same bound). This bound is attained: in a maximal-correlation *MI*-motif (e.g., selected dimers AA, CC, GG, TT; see Appendix B.1), every neighbor of a selected dimer is non-selected, so all four selected dimers are optima.

## Supplementary Material

**Figure S1.**
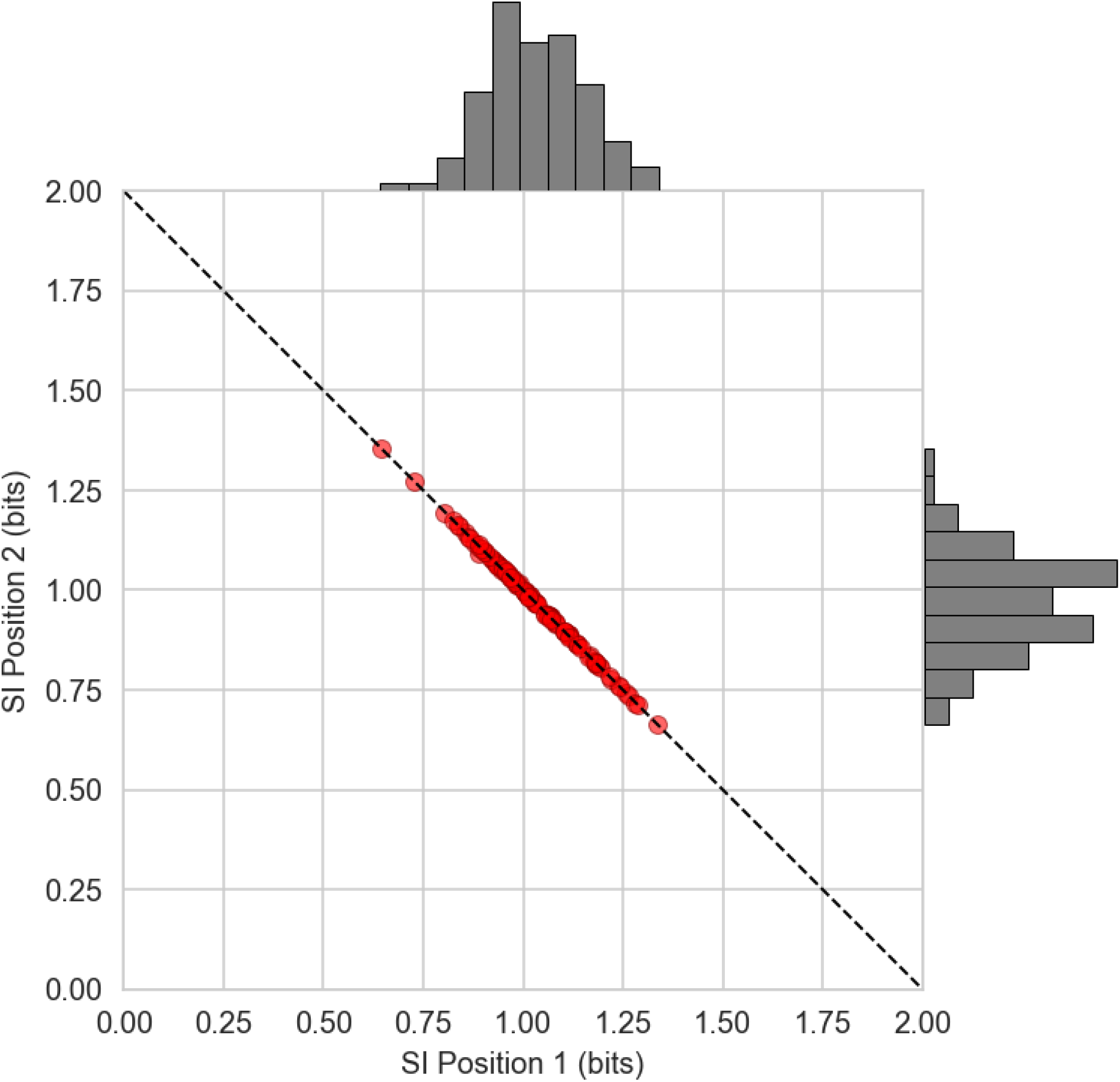
Distribution of single-locus information across motif positions for randomly sampled *SI*-based motifs. Each point represents one of 100 randomly sampled 2 bp-long (*l*_*bs*_ = 2) *SI*-based motifs, consisting of 128 (*n*_*bs*_ = 128) TF-binding sites and encoding 2 bits of total information. The *SI* at position 1 and position 2 are plotted on the horizontal and vertical axes, respectively, with the marginal distributions shown as a histogram. Points cluster near center of the diagonal, indicating that randomly sampled *SI*-based motifs predominantly distribute information approximately equally across positions, consistent with *SI* Case B (Figure 2B). Motifs with strongly asymmetric distributions (*SI* Case A) are comparatively rare.

**Figure S2.**
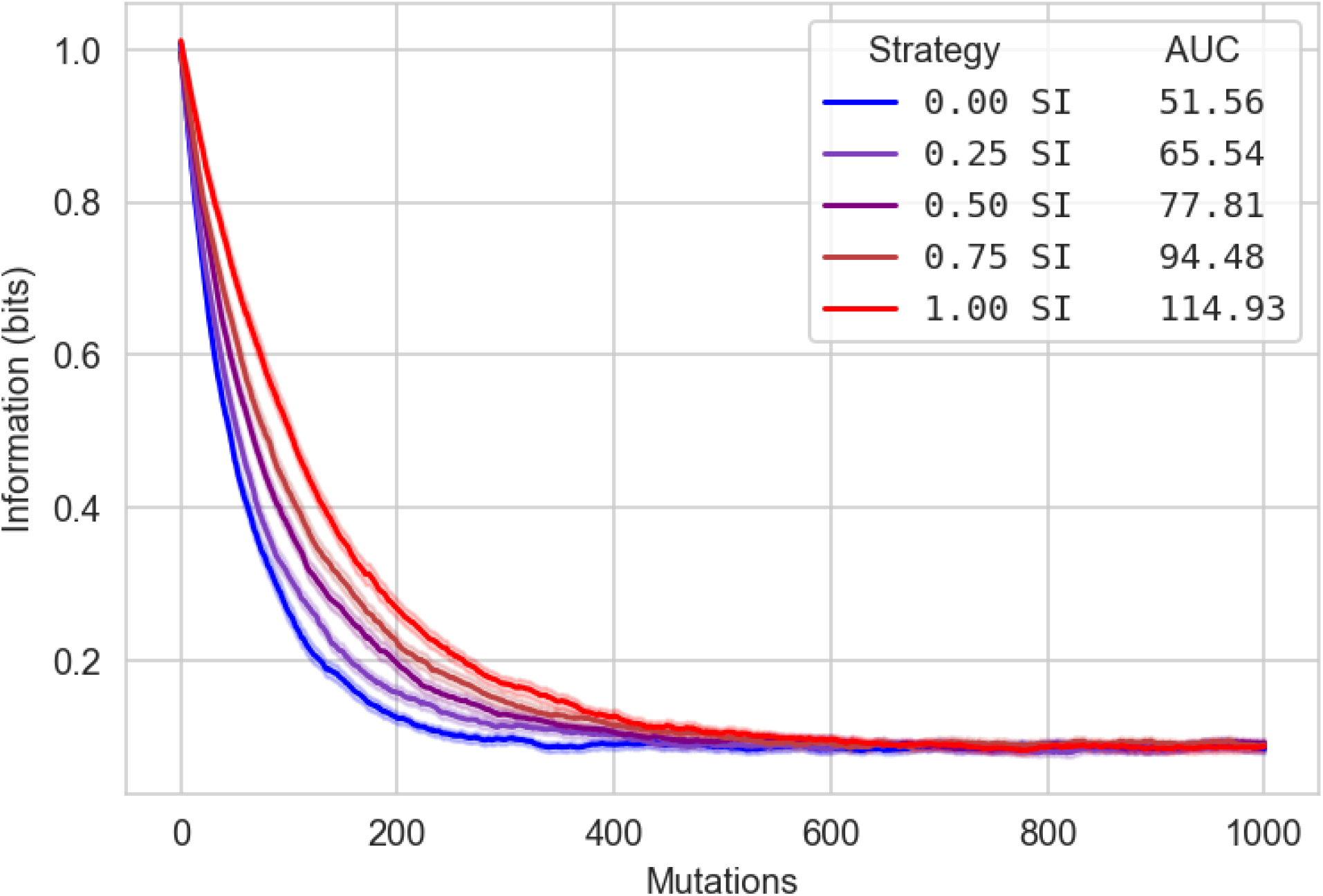
Information loss under mutation for motifs encoding different fractions of single-locus information. Total information (*TI*) of five motif populations as random mutations accumulate on binding sites. Encoding strategies range from 0% *SI* (100% *MI*) to 100% *SI* (0% *MI*) in equal 25% increments. For each strategy, the average (solid line) and the standard error of the mean (shaded area) are plotted for 100 randomly sampled 2 bp-long motifs (*l*_*bs*_ = 2) with 128 binding sites (*n*_*bs*_ = 128). Mutational robustness scales monotonically with the fraction of information encoded as *SI*.

**Figure S3.**
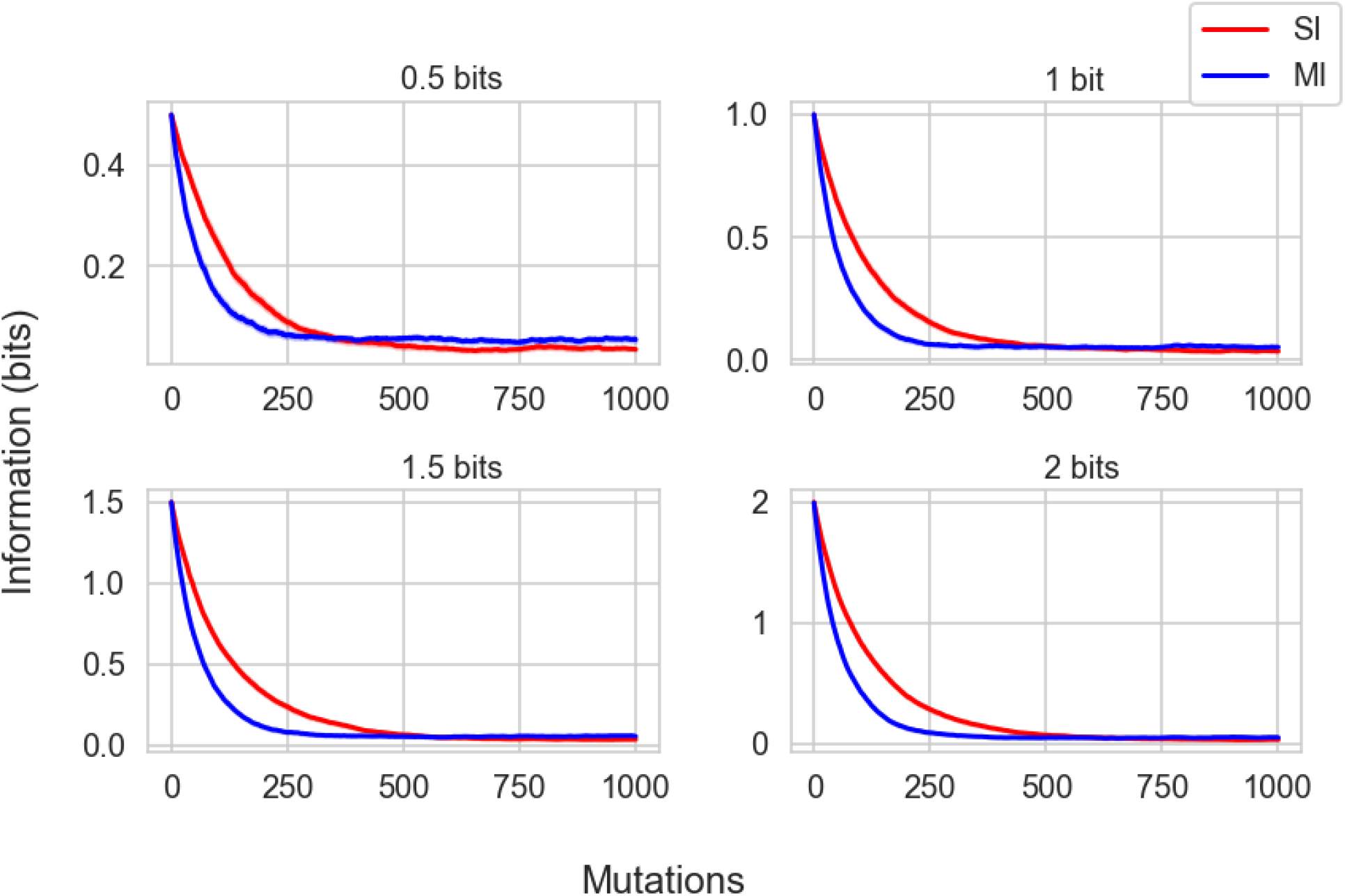
Information loss due to mutation for MI- and SI-based motifs at different information densities. Total information (TI) of SI-based (red) and MI-based (blue) motif populations as random mutations accumulate on binding sites. Information density varies across panels as indicated. For each population, the average (solid line) and the standard error of the mean (shaded area) are plotted for 100 randomly sampled 2 bp-long motifs (*l*_*bs*_ = 2) with 128 binding sites (*n*_*bs*_ = 128).

**Figure S4.**
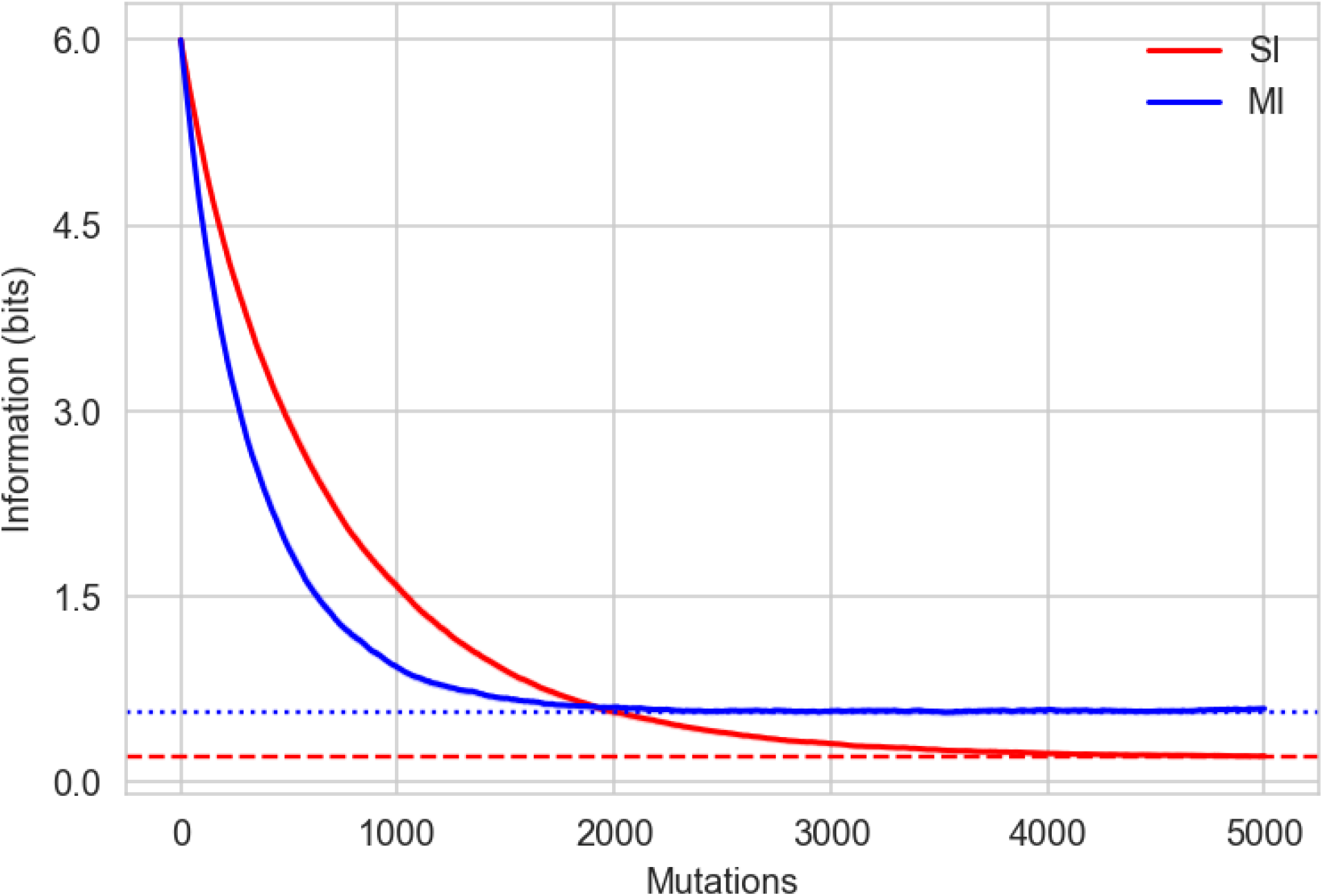
Information loss under mutation for *MI*- and *SI*-based 12 bp-long motif populations. Total information (*TI*) of *SI*-based (red) and *MI*-based (blue) motif populations as random mutations accumulate on binding sites for motifs of length *l*_*bs*_ = 12. The average (solid line) and the standard error of the mean (shaded area) are plotted for 100 randomly sampled motifs with 128 binding sites (*n*_*bs*_ = 128). The red and blue dashed horizontal lines indicate the expected residual *SI* and *MI*, respectively, due to finite sampling effects given the motif depth (*n*_*bs*_ = 128) and length (*l*_*bs*_ = 12).

**Figure S5.**
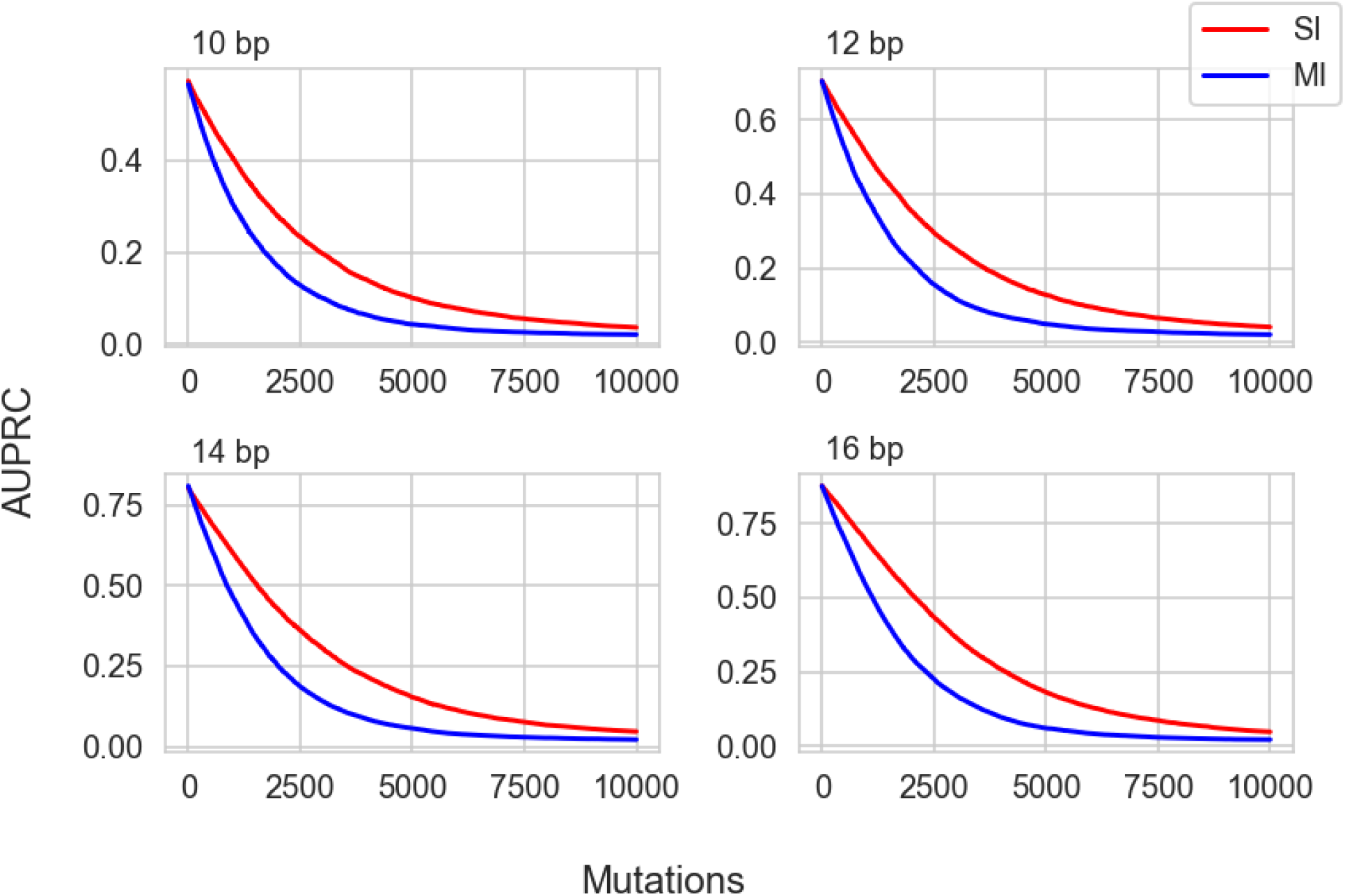
AUPRC decay under mutation for *MI*- and *SI*-based TRN populations across motif lengths. Classification performance, measured as the area under the precision-recall curve (AUPRC), for *SI*-based (red) and *MI*-based (blue) TRN populations as random mutations accumulate. The average (solid line) and the standard error of the mean (shaded area) are plotted for 100 randomly sampled motifs per population. Motif length (*l*_*bs*_) varies across panels as indicated; information density remains constant at *l*_*bs*_/2 bits.

**Figure S6.**
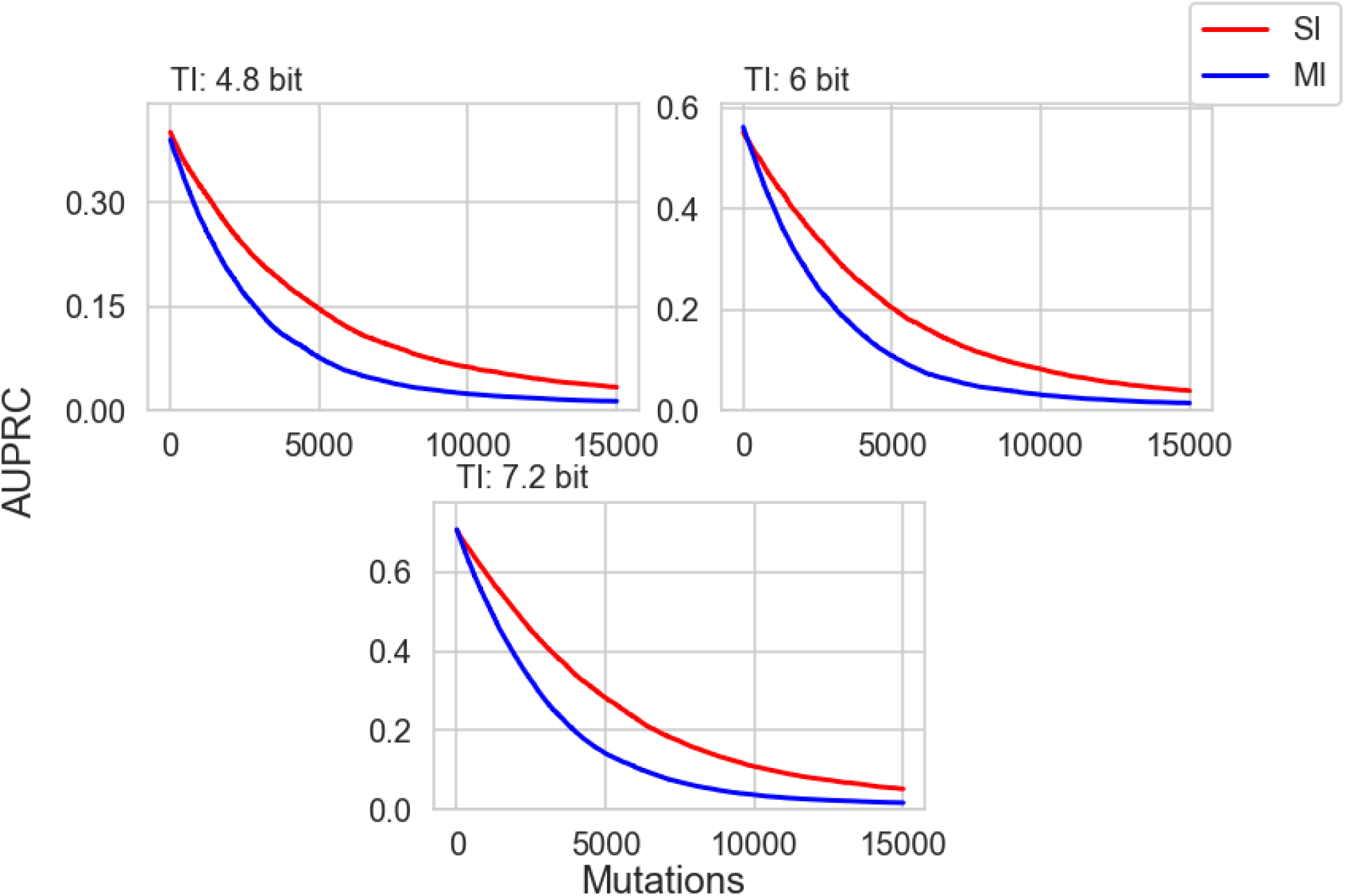
AUPRC decay under mutation for *MI*- and *SI*-based TRN populations across information densities. Classification performance, measured as the area under the precision-recall curve (AUPRC), for *SI*-based (red) and *MI*-based (blue) TRN populations as random mutations accumulate. The average (solid line) and the standard error of the mean (shaded area) are plotted for 100 randomly sampled motifs per population. Total information (*TI*) varies across panels as indicated; motif length remains constant (*l*_*bs*_ = 12).

**Figure S7.**
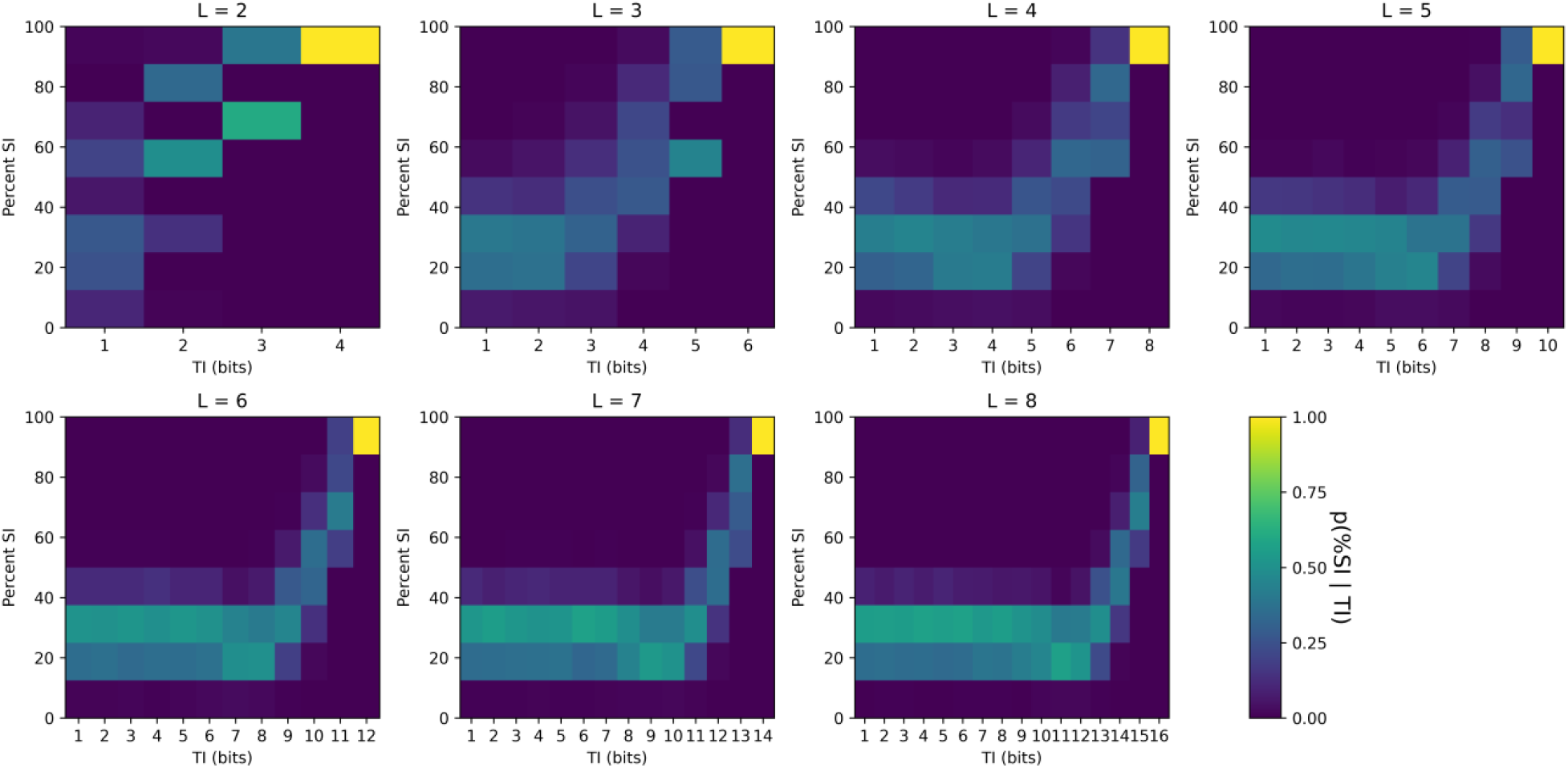
Probability distribution of percent *SI*, conditioned on *TI*, and motif length. Probability distribution of percent *SI* (vertical axis), conditioned on *TI* (horizontal axis), where *TI* is computed as the sum of single-locus information (*SI*) and mutual information among each pair of adjacent positions, for randomly sampled motifs (see Section 4.2) of various lengths (*l*_*bs*_ from 2 to 8 bp). The probability distributions are estimated based on a sample of size 1000.

**Figure S8.**
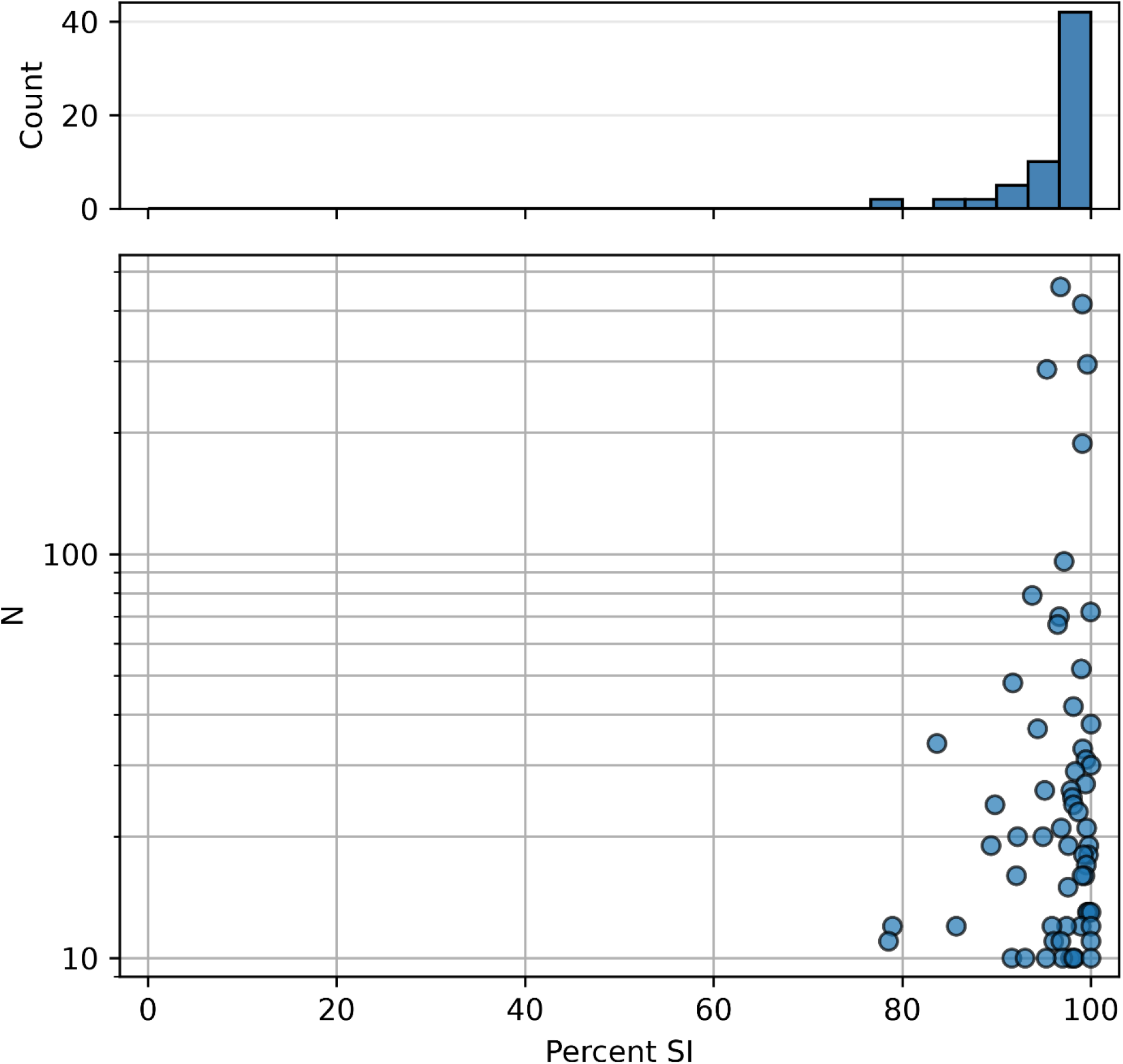
Percent SI in prokaryotic TF motifs. Analysis of the percent *SI* observed in prokaryotic TF-binding motifs from a previously published dataset (https://zenodo.org/records/45329), which was curated in (O’Neill and Erill 2016). The dataset comprises only experimentally validated TF-binding sites. Small-sample corrections are applied to all the observed entropy values. We apply the exact correction (based on the expected value of the entropy under a null uniform sampling) for TF-binding motifs that have up to 90 sites. For TF-binding motifs with a larger number of target sites, we apply the Basharin/Miller-Madow asymptotic approximation (Basharin 1959; G. Miller 1966; G. A. Miller and Madow 1954). The average percent SI for this dataset is 96.35%, with a standard error of 0.60.

**Figure S9.**
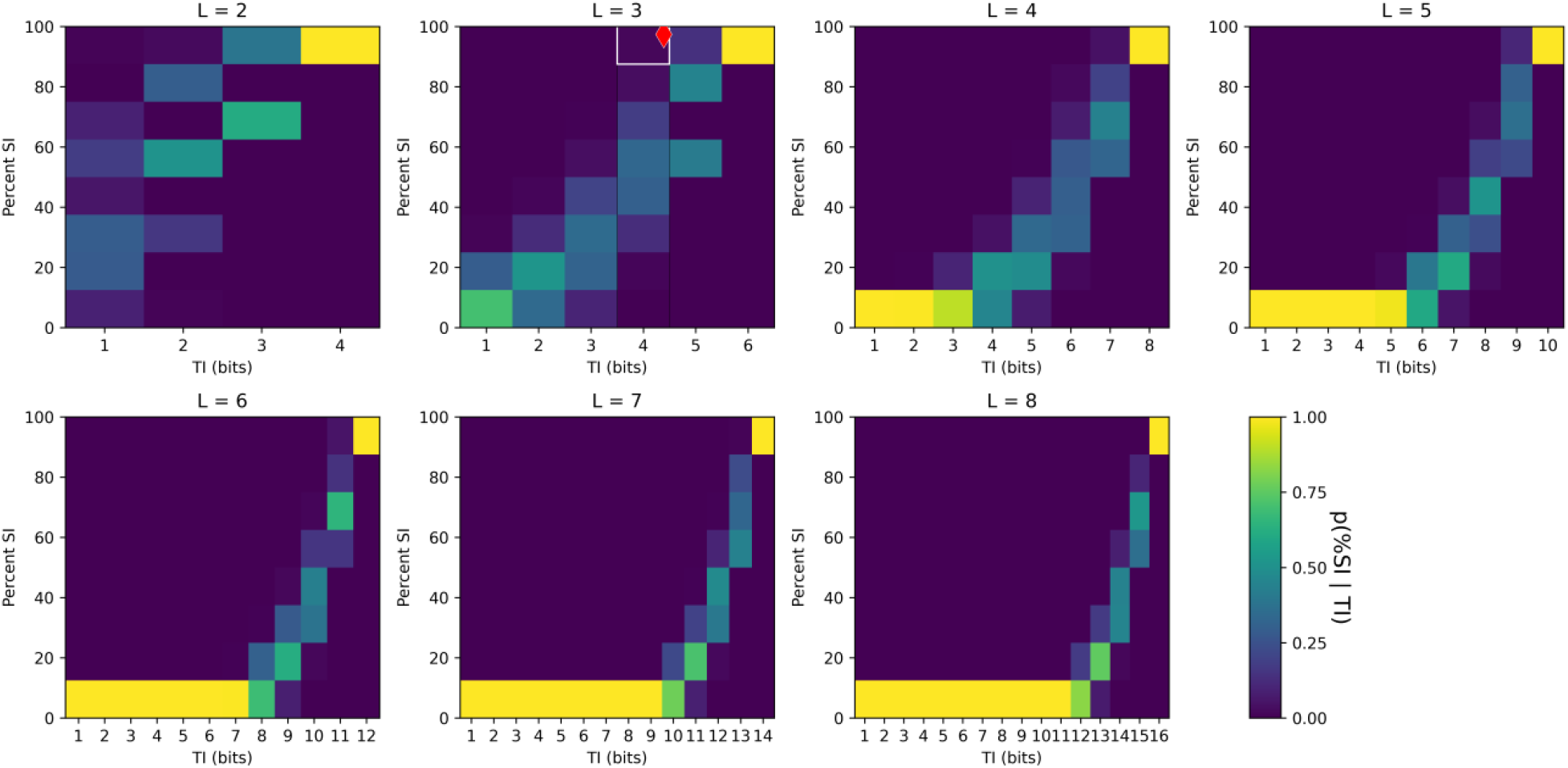
Probability distribution of percent *SI*, conditioned on *TI*, and motif length. Probability distribution of percent *SI* (vertical axis), conditioned on *TI* (horizontal axis), where *TI* is computed as the true total information (Eq. A.7), that is, the sum of single-locus information (*SI*) and the total correlation in the sequence motif (*C*), for randomly sampled motifs (see Section 4.2) of various lengths (*l*_*bs*_ from 2 to 8 bp). The probability distributions are estimated based on a sample of size 1000. The features of the standard genetic code are represented with a red diamond.

**Figure S10.**
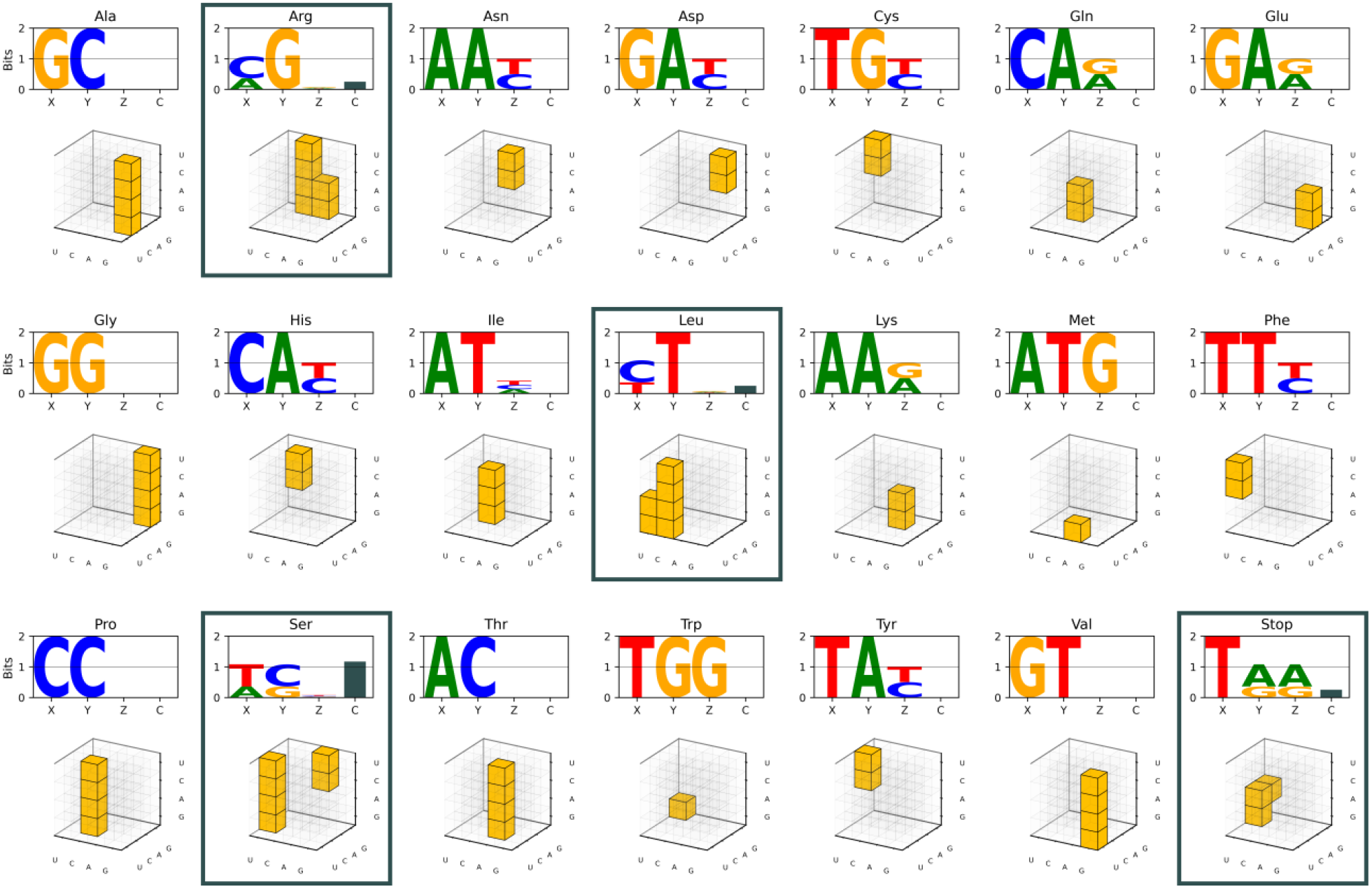
Comparison of sequence logos and genotype tables for the standard genetic code. The 21 standard messages (20 amino acids and one stop signal) are each represented with a sequence logo and as selected genotypes on the (3D) genotype table. Since the codon length is *L*=3, the genotype table appears as a cube. In (Maeshiro and Kimura 1998), the authors noted that when viewed through this cube-based representation, all messages constitute regular subsets that follow specific geometric constraints, except for four messages (bordered boxes). Here, we have added an additional bar to each sequence logo to show the total correlation *C* (in bits) for each message. The four messages with *C* > 0 are precisely the ones violating the above-mentioned geometric constraint (bordered boxes). This relationship is explained in Appendix C.

**Figure S11.**
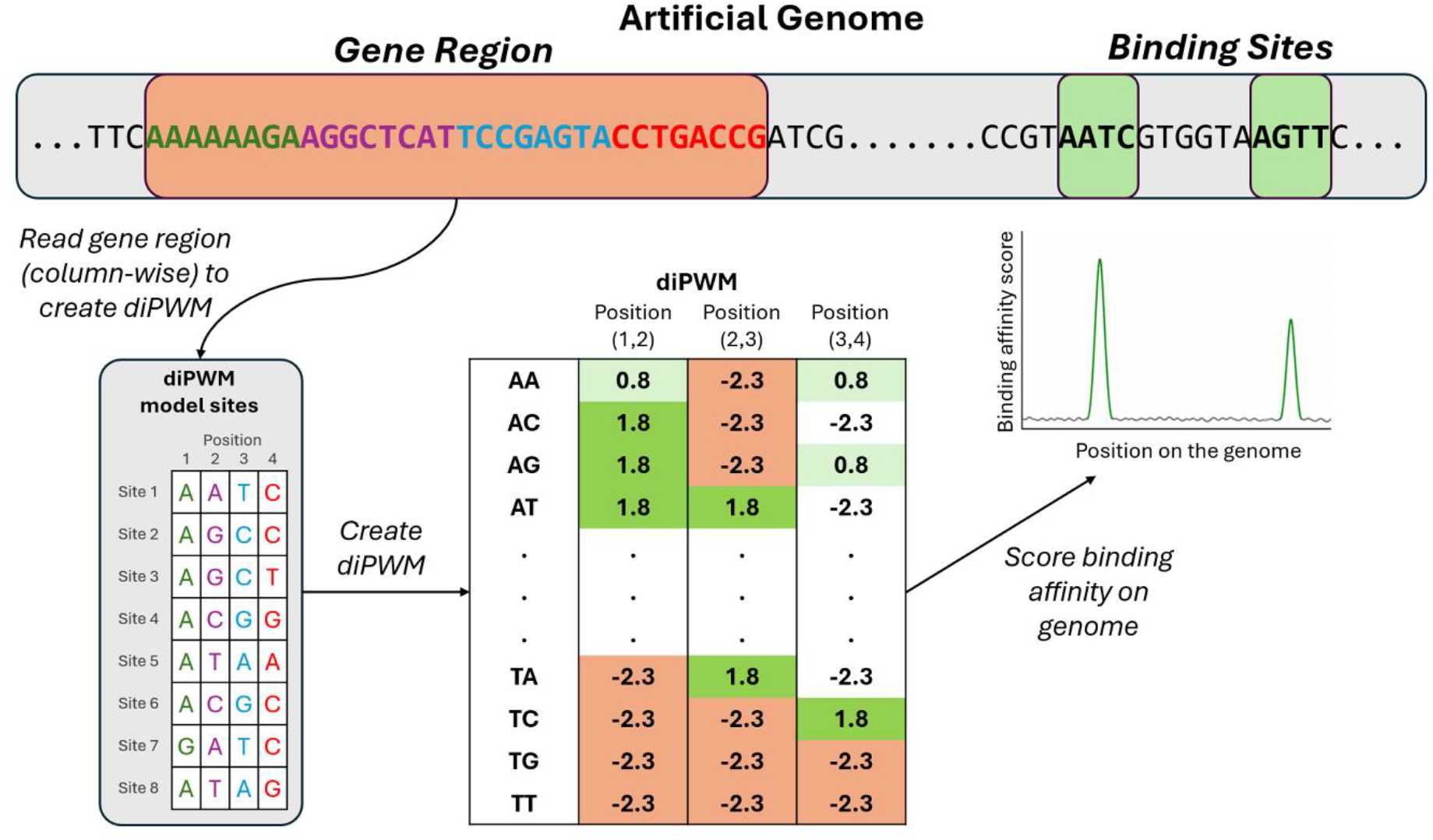
Genome architecture of a simulated organism in the coevolutionary simulations. The genome is a DNA sequence of length *l*_*G,coev*_ = 2048 containing a designated gene region (orange) and pre-designated target sites (green) of length *l*_*bs*_ = 6.The gene region (orange) encodes the TF-binding model as a set of binding sites arranged column-wise, which is read at each generation to instantiate a dinucleotide position weight matrix (diPWM) that captures dependencies between adjacent nucleotide positions. The diPWM model is applied as a sliding window across the genome, producing a vector of binding affinity scores for each genomic position. True target sites (green) yield high scores, while non-target positions (gray) yield low scores.

